# A quantitative model for the dynamics of target recognition and off-target rejection by the CRISPR-Cas Cascade complex

**DOI:** 10.1101/2022.01.26.477710

**Authors:** Marius Rutkauskas, Inga Songailiene, Patrick Irmisch, Felix E. Kemmerich, Tomas Sinkunas, Virginijus Siksnys, Ralf Seidel

## Abstract

CRISPR-Cas effector complexes recognise nucleic acid targets by base pairing with their crRNA which enables easy re-programming of the target specificity in rapidly emerging genome engineering applications. However, undesired recognition of off-targets, that are only partially complementary to the crRNA, occurs frequently and represents a severe limitation of the technique. Off-targeting lacks comprehensive quantitative understanding and prediction. Here, we present a detailed analysis of the target recognition dynamics by the Cascade surveillance complex on a set of mismatched DNA targets using single-molecule supercoiling experiments. We demonstrate that the observed dynamics can be quantitatively modelled as a random walk over the length of the crRNA-DNA hybrid using a minimal set of parameters. The model accurately describes the recognition of targets with single and double mutations providing an important basis for quantitative off-target predictions. Importantly the model intrinsically accounts for observed bias regarding the position and the proximity between mutations and reveals that the seed length for the initiation of target recognition is controlled by DNA supercoiling rather than the Cascade structure.

## INTRODUCTION

CRISPR (clustered regularly interspaced short palindromic repeats)-Cas (CRISPR-associated) systems constitute adaptive RNA-guided defence systems in prokaryotes against foreign nucleic acids (Barrangou et al., 2007; Marraffini, 2015; Wiedenheft et al., 2012). Cas protein effector complexes guided by the crRNA recognize and destroy invading nucleic acids (Koonin et al., 2017; Makarova et al., 2020). Class 2 CRISPR-Cas systems employ single protein effectors as exemplified by Cas9 and Cas12, complexes that were repurposed as genome editing tools in different model organisms from bacteria to human cells (Cong et al., 2013; Jiang et al., 2013; Mali et al., 2013; Zetsche et al., 2017). Effector complexes of the Class 1 systems are arranged of multiple subunits as exemplified by the Cascade complex and recently emerged as a promising tool for genome modification (Cameron et al., 2019; Dolan et al., 2019; Morisaka et al., 2019; Young et al., 2019).

While the effector complexes can be flexibly re-programmed to target practically any unique sequence in a genome (Cong et al., 2013; Mali et al., 2013) there is a significant promiscuity in target recognition that leads to the binding and cleavage of only partially matching DNA sequences (Cho et al., 2014; Cradick et al., 2013; Frock et al., 2015; Fu et al., 2013; Hsu et al., 2013; Kim et al., 2015; Lin et al., 2014; Pattanayak et al., 2013; Tsai et al., 2015; Wang et al., 2015). Such off-targeting can result in highly undesired and unpredictable genetic rearrangements which is particularly problematic for therapeutic applications (Anderson et al., 2018; Cullot et al., 2019; Fu et al., 2013). Off-targeting has been detected using high throughput techniques such as genome-wide *in vivo* DNA binding and cleavage studies (Akcakaya et al., 2018; Fineran et al., 2014; Kim et al., 2015; Tsai et al., 2015, 2017; Wienert et al., 2019), large on-purpose libraries for reporting DNA binding and cleavage *in vivo* (Doench et al., 2016; Hsu et al., 2013; Listgarten et al., 2018) as well as *in vitro* (Boyle et al., 2017; Jung et al., 2017).

The development of engineered effector variants (Amrani et al., 2018; Chen et al., 2017; Edraki et al., 2019; Gleditzsch et al., 2016; Kleinstiver et al., 2016; Lee et al., 2018; Luo et al., 2016; Ran et al., 2015; Slaymaker et al., 2016; Songailiene et al., 2019; Tuminauskaite et al., 2020; Wu et al., 2018) could recently reduce but not abolish off-targeting (Frock et al., 2015; Slaymaker et al., 2016). A frequently used complementary approach to prevent off-targets are *in silico* off-target predictors that promise to identify crRNAs with least promiscuity (Aprilyanto et al., 2021; Bae et al., 2014; Charlier et al., 2021; Haeussler et al., 2016; Lei et al., 2014; Lin and Wong, 2018; Minkenberg et al., 2019; Singh et al., 2015; Stemmer et al., 2015; Xu et al., 2017). Such prediction tools use heuristic scoring functions that try to reproduce sequence and mismatch position patterns from high throughput studies. Though many potential off-target sites are correctly predicted, a considerable fraction of them remains undiscovered by such algorithms (Kim et al., 2015; Newton et al., 2019; Tsai et al., 2015, 2017), such that off-targeting persists to be a challenging problem of CRISPR-Cas technologies.

Along with extensive characterization of off-targeting, considerable mechanistic insight into the target recognition process by CRISPR-Cas effector complexes has been obtained from biochemical, structural and single-molecule studies. A converging theme emerged (Figure 1A) in which an effector complex first scans duplex DNA for a short complex-specific protospacer adjacent motif (PAM). Upon PAM recognition, it initiates base pairing between the crRNA and the PAM adjacent bases of the DNA target strand. The RNA-DNA heteroduplex can then reversibly expand expelling the non-target DNA strand and forming a triple-stranded R-loop structure. Upon full-length R-loop formation up to the PAM-distal end, a conformational change occurs that licenses DNA degradation. For Cascade this comprises a global sliding of the Cse1-Cse2 filament (Wiedenheft et al., 2011; Xiao et al., 2017) that locks the R-loop in a highly stable conformation (Rutkauskas et al., 2015; Szczelkun et al., 2014) and allows recruitment of the Cas3 nuclease (step iv in Figure 1A). The actual target recognition is, however, a strand displacement reaction between the involved nucleic acid strands (steps ii and iii in Figure 1A), in which the effector complex acts only as a sensor for the R-loop progression. Mismatches can be considered as energy barriers during the reversible R-loop expansion that promote its collapse (Rutkauskas et al., 2015). Notably, the mismatch strength has shown to be biased with respect to the position; PAM-proximal mismatches within the so-called seed region have been shown to impose stronger inhibition of R-loop formation compared to distal mismatches (Fineran et al., 2014; Semenova et al., 2011).

**Figure 1.**
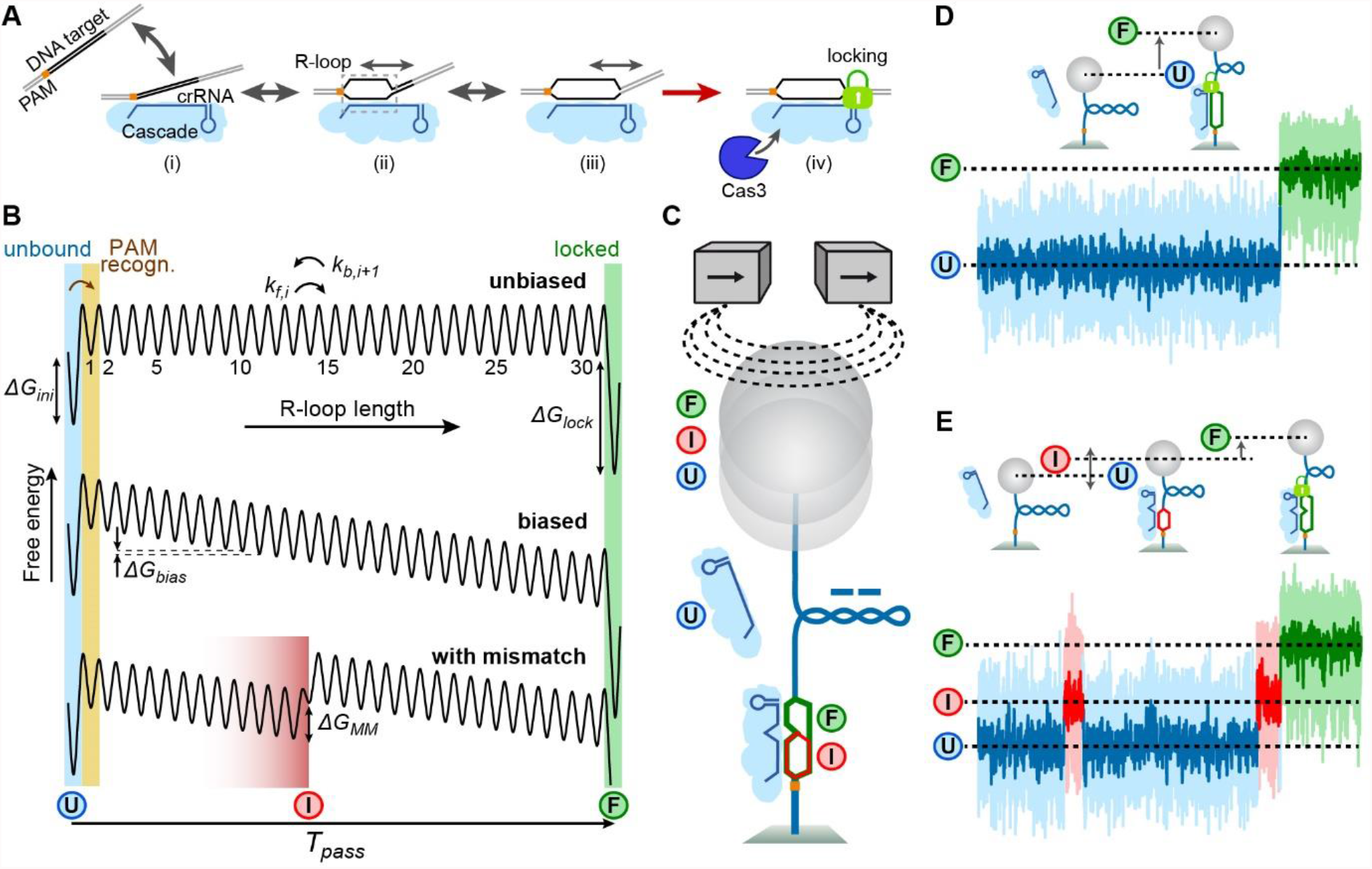
Random-walk model for target recognition by Cascade. (A) Target recognition starts with PAM binding (i) followed by R-loop priming adjacent to the PAM. The R-loop can expand in a reversible random-walk like fashion towards the PAM distal end of the target (ii). After reaching PAM distal end (iii), a conformational change is induced that locks the R-loop in a stable conformation and enables Cas3 recruitment and cleavage of the DNA (iv). (B) Simplified one-dimensional energy landscapes for R-loop formation. Local energy states are the unbound state (*U*), the R-loop states of different length and the locked full R-loop (*F*). R-loop expansion and shrinkage occurs in successive forward or backward steps to either adjacent state. Shown energy landscapes correspond to R-loop formation without an energy bias (top), with a negative bias as expected for negative supercoiling or due to favorable molecular interactions (middle) and with a mismatch at position 14, which introduces a local energy barrier (bottom). In the latter case an additional apparent intermediate R-loop state (*I*) is introduced. (C) Real-time detection of the R-loop dynamics on negatively supercoiled DNA using magnetic tweezers. R-loop expansion progressively absorbs part of the introduced supercoiling, which results in a DNA length increase. (D) R-loop formation on a fully matching target is seen as a single abrupt DNA length increase corresponding to the transition from the unbound to the full-R-loop state. (E) R-loop formation on a target with a single internal C:C mismatch at position 17 occurs via additional transitions to and from the unstable intermediate state until the full locked R-loop is formed.

Despite an increasing and detailed mechanistic understanding of target recognition by CRISPR-Cas complexes, the wealth of mechanistic knowledge has been barely linked to the heuristic scoring schemes used in off-target predictions. Importantly the reversible nature of the R-loop expansion together with the irreversible locking or cleavage steps impose a new kinetic rather than a “sticky”, i.e. affinity-based, target recognition mechanism (Bisaria et al., 2017) which is governed by the competition between R-loop collapse and expansion. Such a kinetic scheme has, however, not been applied in scoring systems nor, despite being widely suggested been quantitatively applied to the targeting dynamics in mechanistic studies (Bisaria et al., 2017; Klein et al., 2018; Singh et al., 2018).

Here we use single-molecule DNA twist measurements to comprehensively quantify the dynamics of R-loop formation by the Cascade complex from *S. thermophilus*. Importantly, we resolve the transient R-loop sub-states on single- and double-mismatched DNA targets which remained hidden for other methods including high-throughput off-targeting measurements (Boyle et al., 2017; Jung et al., 2017). We show that the observed dynamics can be quantitatively described by a random walk model in a simplified one-dimensional energy landscape. The model was adapted from previous descriptions of protein-free strand displacement reactions in dynamic DNA nanotechnology (Irmisch et al., 2020; Machinek et al., 2014; Srinivas et al., 2013), which have recently also been introduced to the CRISPR-Cas field (Eslami-Mossallam et al., 2020; Klein et al., 2018). Importantly, our modelling (i) provides direct evidence that R-loop expansion follows a random-walk process (ii), shows that the single-base pair stepping of R-loop expansion occurs at a sub-millisecond time scale, (iii) returns absolute free energy penalties imposed by mismatches, (iv) explains the non-trivial dependence of R-loop formation on the proximity between multiple mismatches and (v) reveals that the length of the seed region in Cascade is a function of the applied supercoiling rather than a structural property.

Overall, our findings establish an important mechanism-based approach for the prediction of off-targeting by Cascade and potentially other genome engineering tools. We furthermore note that our model directly includes the dependence of off-targeting on DNA supercoiling, which in eukaryotes has been found to be highly locus- and gene specific (Corless and Gilbert, 2017; Kouzine et al., 2013; Naughton et al., 2013).

## RESULTS

### Modeling R-loop formation as random-walk on a one-dimensional energy landscape

Previous single molecule experiments of target recognition by Cascade (Krivoy et al., 2018; Rutkauskas et al., 2015) strongly suggested that following binding to the AAN PAM (Table S3), Cascade nucleates an R-loop at the PAM proximal end of the target sequence. The R-loop can then further expand towards the PAM distal end in a fully reversible manner until locking occurs (Figure 1A). Mismatches were found to be barriers for R-loop expansion such that they promoted R-loop collapse. Based on this model as well as on previous understanding of DNA strand displacement reactions (Irmisch et al., 2020; Liu et al., 2021; Machinek et al., 2014; Srinivas et al., 2013), we hypothesize that an R-loop is a dynamic structure which grows and shrinks in single base-pair steps in a random-walk-like fashion (Josephs et al., 2015; Klein et al., 2018). To build a quantitative model, as first approximation, we constructed a simple one-dimensional energy landscape for R-loop formation with the R-loop length as reaction coordinate. Cascade starts initially in the unbound state (*U*) corresponding to an R-loop of zero length. Upon PAM binding an R-loop of 1 bp length shall be nucleated for which a free energy penalty *ΔG*_*ini*_ is considered. In a first iteration we assumed that the free energy for increasing R-loop lengths is constant, since for each additional base-pair of the heteroduplex a base pair of the DNA-duplex needs to be disrupted (Figure 1B, top). After full R-loop formation of 32 bp for Cascade, the R-loop enters the locked state (state *F*) associated with a decrease in free energy by *ΔG*_*lock*_. Negative supercoiling has been shown to assist R-loop formation (Ivanov et al., 2020; Rutkauskas et al., 2015; Szczelkun et al., 2014). This enters as a constant negative bias *ΔG*_*bias*_ for increasing R-loop lengths (Figure 1B, middle). Also, protein contacts to the R-loop can contribute an additional favorable or unfavorable bias that we assume to be constant throughout the R-loop. When overcoming a mismatch, all following states are shifted in the positive free energy direction by the mismatch penalty *ΔG*_*MM*_ (Figure 1B, bottom). Thus, a mismatch would introduce a dynamic intermediate R-loop state *I*, which extends over a few base pairs before the mismatch due to the random-walk nature of the R-loop (Figure 1B, bottom). For *ΔG*_*bias*_ = 0, we assume the kinetic barriers for all transitions between R-loop states to be identical except for transitions from the *U* and *F* states. *k*_*step*_ shall be the unbiased single base-pair stepping rate for conventional R-loop expansion/retraction steps. Rates for transitions with altered energetic barriers including changes due to a non-zero bias and mismatches are then provided by the corresponding free energy penalties according to Arrhenius’ law (see Methods for details). This way one obtains a fully parameterized linear rate model. Mean transition times between states can be obtained by solving the first passage problem for this model (see Supplementary Note 1). This allows to calculate the mean R-loop formation time but also rates for transitions between different intermediate states using the only free parameters *ΔG*_*ini*_, *ΔG*_*bias*_, *k*_*step*_ and in case of mismatches one or several different values for *ΔG*_*MM*_.

### Resolving and quantifying the dynamics of R-loop intermediates

To test the applicability of the random walk model, we set out to quantify the R-loop dynamics of Cascade using single-molecule DNA twisting experiments (Krivoy et al., 2018; Rutkauskas et al., 2015; Szczelkun et al., 2014). Surface-grafted DNA molecules tethering a magnetic bead on their free end were stretched vertically in a magnetic tweezers apparatus (Kemmerich et al., 2016; Rutkauskas et al., 2017). Negative supercoiling introduced by rotating the tweezers magnets provided a DNA length reduction due to formation of writhe (Figure 1C, (Brutzer et al., 2010; Rutkauskas et al., 2015; Strick et al., 1996)). DNA unwinding due to R-loop formation by Cascade absorbs part of the introduced supercoiling causing a DNA length increase (Figure 1C and D, (Rutkauskas et al., 2017)). On a fully matching target this is seen as a single abrupt transition from the unbound (*U*) to the full (*F*) R-loop state (Figure 1D). In the presence of a single internal mismatch, three distinct states are observed – initially, the R-loop reversibly transits between the *U* state and the intermediate state *I* before the mismatch. Upon mismatch passage, it can finally reach the *F* state (Figure 1E). Unlike fluorescence-based approaches (Mekler et al., 2017; Singh et al., 2018; Sternberg et al., 2014), the DNA twisting experiments allow a direct quantification of the R-loop length from the measured length changes (Szczelkun et al., 2014) and this way to unambiguously resolve and characterize the dynamics of the different R-loop states.

### Random walk describes dwell of intermediate R-loops

We first quantified the dwells of intermediate R-loops of different length. The targets contained a limited number of matching base pairs adjacent to the PAM (from 8 to 22 bp) with the remaining base pairs being mismatched. This way only transitions between the *U* and the *I* states were observed (Figures 2A and S1A). We furthermore applied different negative supercoiling levels, given as mechanical torque *τ*, that were controlled by the applied stretching forces (see Methods). Natural superhelical densities *σ* in *E. coli* cells are in the range of −0.06 and −0.029 corresponding to torques *τ* between −8.9 and −4.3 pN nm (Methods, (Brouns et al., 2018; Higgins and Vologodskii, 2015)). Qualitative inspection of the obtained trajectories revealed that the dwell in the *I* state increases with increasing negative supercoiling (Figure 2B) and increasing length of the R-loop intermediate (Figure 2C). This intuitively agrees with the energy landscape of the random-walk model, since increased bias and length lower the free energy of the *I* state and thus increase the energy barrier for a diffusive return to the *U* state (Figure 2F). This is also seen in increased occupancies when modelling the *I* state (Figure 2F). Quantitative analysis of the dwells in both states provided the R-loop formation rates *k*_1_ (Figure 2D) and collapse rates *k*_2_ of the *I* state (Figure 2E). *k*_1_ was rather independent of the R-loop length and the applied supercoiling. This agrees with a short distance of the transition barrier from the *U* state (approximately at the transition to position 1). In contrast, the R-loop collapse rate *k*_2_ was strongly torque- and R-loop length-dependent and varied over three orders of magnitude. A global fit of the random-walk model to all collapse rates correctly described both the large spread of the rates between the different R-loop lengths as well as the torque dependence for a given R-loop length. This provides strong evidence that R-loop expansion and retraction occurs via the proposed random-walk mechanism. Remarkably, fitting only required only one free parameter, i.e. the unbiased single base-pair stepping rate for which we obtained *k*_*step*_ = 1900 ± 100 *s*^−1^. Single base-pair steps during R-loop formation thus occur on the sub-millisecond time scale. Beyond the rates, also the occupancies of the *U* and *I* states were correctly described including their torque and length dependencies (see histograms in Figures 2A and 2B) in line with the intuitive understanding of the minima in the free energy landscape being occupied according to Boltzmann statistics (Figure 2F).

**Figure 2.**
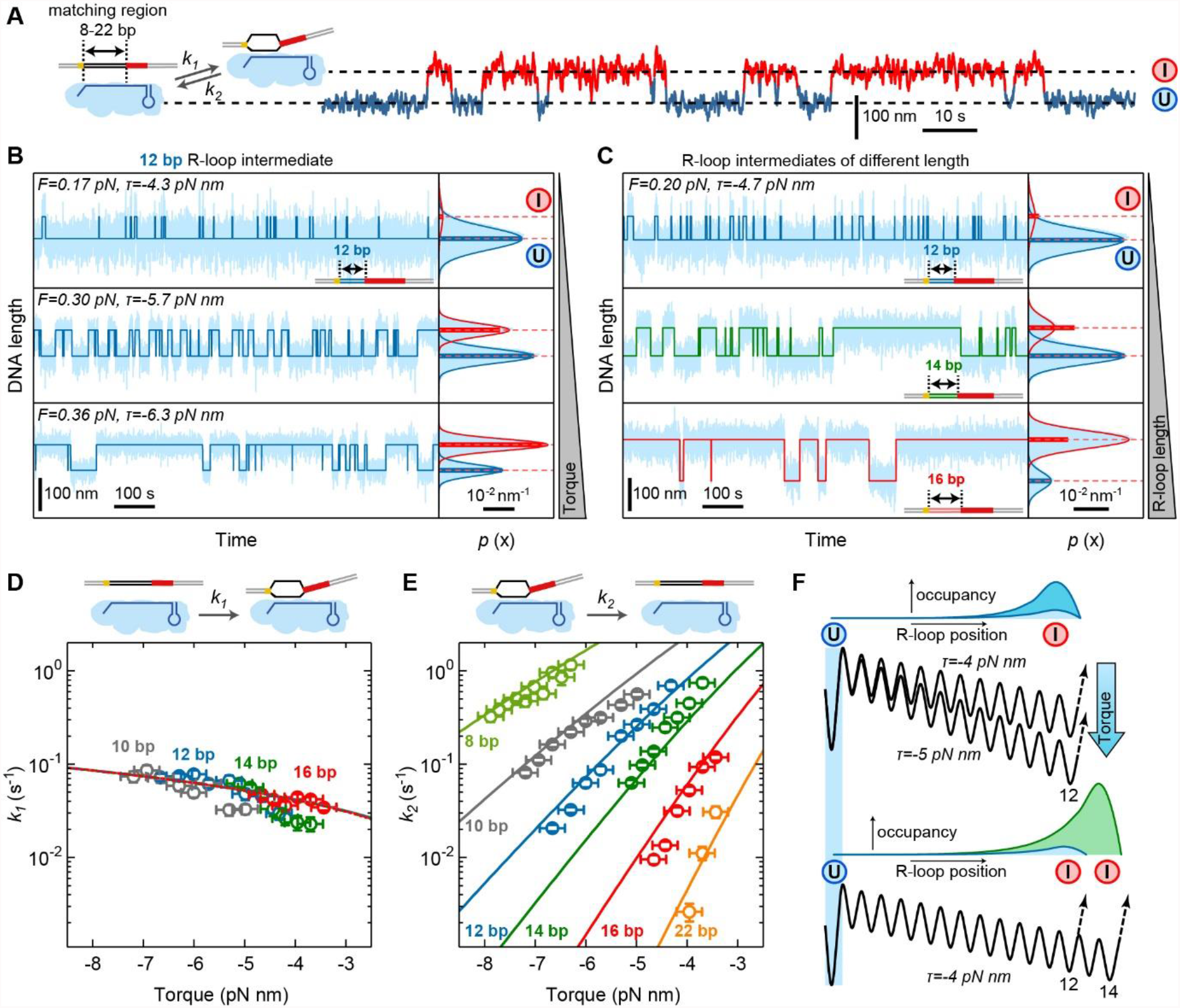
Formation and collapse rates of R-loop intermediates of different length. (A) Sketch of the DNA construct containing a limited number of matching PAM-proximal bases such that only intermediate R-loop intermediates form (left). Example trajectory of the transitions between unbound state (*U*) and intermediate R-loop (*I*) for a target with 12 matching bases. (B) Formation and collapse of R-loop intermediates recorded at different torques *τ* for a target with 12 matching bases. Shown are the recorded DNA length smoothed to 7.5 Hz (light blue) and a two-state approximation of the trajectory (dark blue). Solid lines in the histograms on the right represent Gaussian fits to the 2 different states, while horizontal dashed lines indicate the average DNA length of each state. Bars represent theoretically predicted occupancies using best fit parameters from (D) and (E). (C) Formation and collapse of R-loop intermediates of different length at a torque of -4.7 pN nm. (D,E) Measured mean formation (D) and collapse (E) rates of the R-loop intermediates as function of torque for different lengths of the matching region (open circles). A global fit to the data is shown as solid lines. Rate errors are given as SEM; torque errors correspond to 0.25 pN nm. (F) Occupancies of the *I* state for two different negative torques (upper panel) and R-loop lengths (lower panel) together with the corresponding energy landscapes.

### R-loop dynamics at mismatched targets provides mismatch penalties

We next studied the R-loop dynamics at single target mutations. A single mutation was introduced within a target possessing six additional consecutive mismatches at the PAM-distal end to prevent R-loop locking (Rutkauskas et al., 2015; Szczelkun et al., 2014). On such a target, the R-loop fluctuated between three states – the *U*, the *I* and a dynamic *F*^*\**^ state of a maximum length of 26 bp (Figure 3A). Testing the three possible mismatches C:C, C:T and C:A at position 17 (counting from the PAM) revealed that the mismatch type strongly influenced the transition rates and the occupancies of the three states (Figure 3B). Furthermore, these parameters were influenced by increasing negative torque, where the *F*^*\**^ state became increasingly populated at the expense of the *U* state (Figure 3C). We quantitatively analyzed the four rates that describe the transitions between adjacent states (Methods, Figure S3). This revealed that the rates and their torque dependences were largely independent of the mismatch type except of *k*_3_ that described the mismatch passage from the *I* to the *F*^*\**^ state (Figures 3D and 3E). Generally, rates describing R-loop expansion (*k*_1_, *k*_3_) increased with increasing negative torque, while rates describing R-loop retraction (*k*_2_, *k*_4_) were found to decrease. For a given mismatch, we applied a global fit to the torque dependence of all four rates (solid lines in Figures 3D and 3E). This yielded good agreement with the data supporting the applicability of our model. The best fit parameters for *k*_*step*_ (Figure 3F) and *ΔG*_*ini*_ (Figure 3G) were mismatch independent while the mismatch penalty *ΔG*_*MM*_ (Figure 3H left panel, Table S1) was strongly mismatch dependent. This is consistent with the intuitive expectation that the mismatch type influences only the corresponding penalty but not the other parameters. Applying different Cascade concentrations revealed a linear dependence of the initial R-loop intermediate formation rate *k*_1_ on the concentration while leaving the other rates unchanged (Figure S2) in agreement with concentration-independent values for the standard free energy of R-loop initiation and the other model parameters (Figure S2C). As an independent test of our model, we used the best fit parameters describing the R-loop dynamics to calculate the expected occupancies of the *U, I* and *F*^*\**^ states and observed good agreement with the experimental data (Figures 3B and 3C right panels). Altogether, the random walk model could successfully describe the 3-state dynamics over a single mismatch.

**Figure 3.**
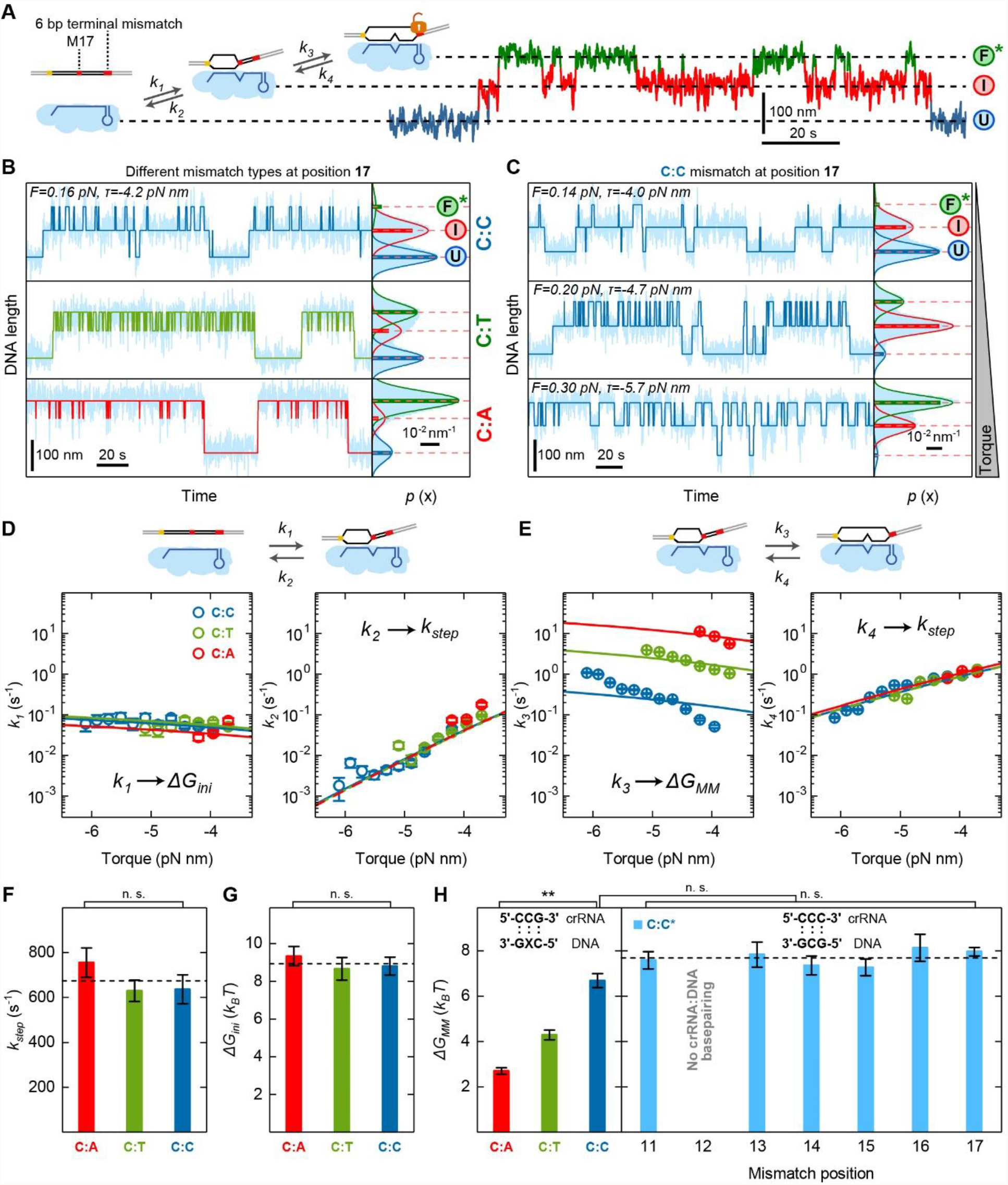
R-loop dynamics on locking-deficient targets containing a single internal mismatch. (A) Sketch of the DNA construct with mismatched bases shown in red and the possible R-loop states - the unbound state (*U*), the intermediate R-loop (*I*) and the almost full but unlocked R-loop (*F*^*\**^). Sequential transitions between these states can be described by 4 different rate constants (*k*_1_ to *k*_4_) as indicated. Example trajectory of the transitions between the different states measured for a C:C mismatch. (B) Trajectories and histograms of the DNA length recorded for targets with either a C:C, C:A or a C:T mismatch at a torque of 4.2 pN nm (light blue). 3-state approximations of the trajectories in B and C are shown in dark blue (C:C mismatch), green (C:T mismatch) or red (C:T mismatch). (C) Trajectories and histograms of the DNA length recorded for different applied torques using a target with C:C mismatch at position 17 (light blue). Solid lines in the histograms represent Gaussian fits to the 3 different states, while horizontal dashed lines indicate the average DNA length of each state. Bars represent theoretically predicted occupancies using the parameters shown in SI Table1. (D) Experimental transition rates between the *U* and the *I* states as function of torque for C:C, C:T and C:A mismatches shown as blue green and red circles, respectively. (E) Experimental transition rates between the *I* and the *F*^*\**^ state as a function of torque. The solid lines shown in D and E correspond to a global fit to all four transition rates for a given mismatch. (F) Single base-pair stepping rates for the different mismatches obtained from the transition rate fits. The dashed line represents the mean rate. (G) Free energy change of R-loop initiation at a Cascade concentration of 170 nM for the different mismatches obtained from the transition rate fits. The dashed line represents the mean. (H) Mismatch penalties for the different mismatches at position 17 (left panel) and of the C:C mismatch at different positions (right panel) obtained from the transition rate fits. Error bars in all plots correspond to SEM. n. s. – no significance; **p<0.01.

Next, we investigated how *ΔG*_*MM*_ depended on the mismatch position. When keeping the same mismatch type including the same nearest-neighbor base pairs, *ΔG*_*MM*_ should be largely position independent if nucleic acid thermodynamics would dominate. To test this, we produced different Cascade complexes with a CCC stretch at different positions in the crRNA (Table S3 and S4) allowing a corresponding introduction of a C:C mismatch with two adjacent G:C base pairs (see sketch in Figure 3H, right panel). Recording and fitting the torque dependence of the different transition rates for these complexes revealed that *ΔG*_*MM*_ was within error invariant for mismatch positions from 11 to 17 bp that could be experimentally accessed (Figure 3H right panel, Table S1). Note that mismatch barriers were not observable at positions 6, 12, 18, 24 and 30 (Fineran et al., 2014; Rutkauskas et al., 2015) due to the disrupted base pairing in the crRNA-DNA hybrid at these positions (Mulepati et al., 2014).

### Seed length is dependent on supercoiling

A mismatch penalty that is rather independent of the mismatch position seems to apparently contradict the larger impact of PAM-proximal mismatches in the seed region compared to PAM-distal mismatches observed *in vivo* (Fineran et al., 2014; Semenova et al., 2011) and the measurements of apparent mismatch penalties in high-throughput experiments (Jung et al., 2017). We therefore tested whether our model would describe such a position-dependent inhibition of R-loop formation correctly, despite a uniform mismatch penalty. To this end, we expanded the range of mismatch positions (positions 5 to 21). Since for many of these targets intermediates were too short lived to be observable, we measured only the time of the R-loop formation (state *F*) using a full-length target that supported locking (Figures 4A and S1B-S1E). In agreement with previous reports (Rutkauskas et al., 2015; Semenova et al., 2011), R-loop formation was slower for targets with PAM-proximal mismatches compared to PAM distal mismatches and the WT target (Figures 4B, S4A and S4B). We determined the mean R-loop formation time as a function of torque for the WT target and the single mismatch targets (Figures 4C, S4A and S4B). R-loop formation for the WT target was little dependent on torque in the applied range. The R-loop formation times for single-mismatch targets decreased, however, strongly in a non-linear fashion with increasing negative torque and finally plateaued at the WT level (Figures 4D and S4D). The torque required for reaching the WT level was changing monotonously with mismatch position. We applied a global fit of the random walk model to the data using *ΔG*_*ini*_ = 8.5 *k*_*b*_*T* and obtained agreement with the experimental results (Figures 4D and S4D). The consideration of single mismatch targets alongside the WT target in this data set as well as the highly non-linear torque dependence allowed furthermore to probe a potential bias *ΔG*_*bias*_ of the energy landscape in absence of torque. Global fitting with a free non-zero bias provided as best fit parameters *ΔG*_*MM*_ = 6.7 ± 0.4 *k*_*b*_*T, k*_*step*_ = 1600 ± 600 *s*^−1^ as well as *ΔG*_*bias*_ = 0.13 ± 0.01 *k*_*b*_*T*/*bp*. The positive value of *ΔG*_*bias*_ reveals that in absence of torque the energy landscape has a small upward bias (Figure S4E) corresponding to an apparent torque of ∼1 pN nm which is much smaller than base pairing energies and mismatch penalties. Inclusion of the determined *ΔG*_*bias*_ into fits of the previous data were mainly compensated by changes of *k*_*step*_ while the obtained free energy values became only slightly reduced (see Table S2).

**Figure 4.**
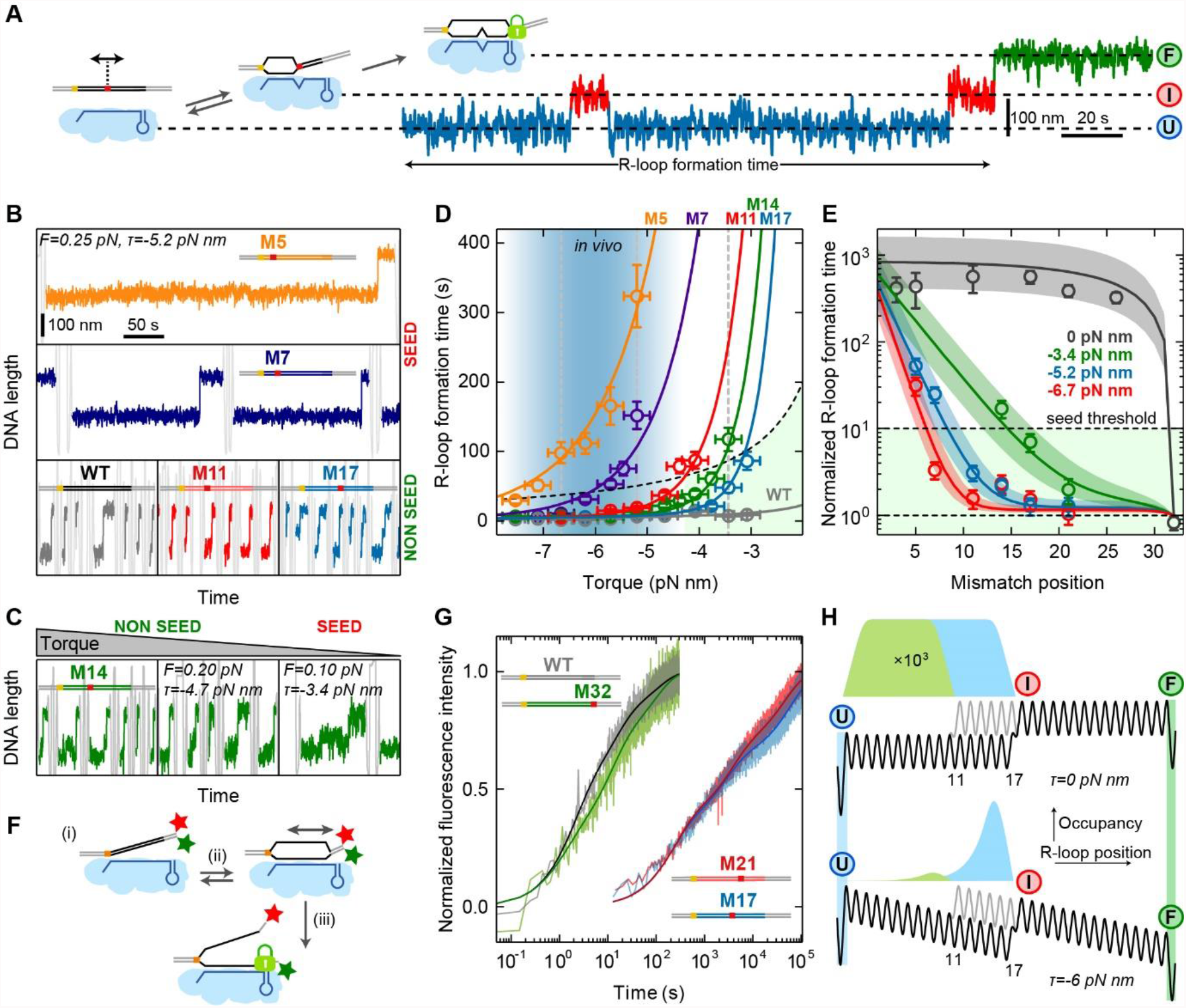
Locked R-loop formation on targets with single internal mismatches. (A) Sketch of the DNA construct containing a single C:C mismatch at various positions along the target (left). Example trajectory of full R-loop formation for a mismatch at position 17 after applying negative supercoiling. The full R-loop is formed after multiple intermediate state events. (B) Successive R-loop formation events for the WT target (gray) and targets with mismatches at position 5 (orange), 7 (purple), 11 (red) and 17 (dark blue) at a torque of 5.2 pN nm. Colored sections represent the actual R-loop formation events, while gray sections correspond to changes of the DNA supercoiling. The length of a formation event is roughly proportional to the R-loop formation time, such that a visual impression of the R-loop formation time for the different mismatches and torques can be obtained. Trajectories represent data smoothed to 3 Hz. (C) Successive R-loop formation events measured at different torques for the target containing mismatch at the position 14. Trajectories are represented as in B. Force and torque values in the first panel are as in B. (D) Torque dependence of the R-loop formation time for different mismatch positions. Circles represent mean times from >30 events collected on at least two molecules. Solid lines represent a global fit of the random-walk model to the data. The light blue gradient marks the torque range estimated *in vivo* (see Methods). The light green area corresponds to a less than 10-fold increase of the R-loop formation time compared to the WT. Errors of the formation time are SEM; and of the torque 0.25 pN nm. (E) Normalized R-loop formation time with respect to the WT (open circles, obtained from data in C, D as well as G) as function of the mismatch position for three different torques (corresponding to the vertical dashed lines in C). Solid lines show the predictions of the random-walk model using the fit parameters from C. Colored areas represent confidence intervals of the fits. (F) Scheme of the fluorescence bulk solution measurements of the R-loop formation kinetics in absence of supercoiling (zero torque) involving a donor-acceptor dye pair at the PAM distal end of the target. The donor fluorescent signal increases mainly due to R-loop formation and locking (ii, iii) but not initial PAM binding (i). (G) Kinetics of R-loop formation in absence of supercoiling for WT and mismatched targets. (H) Comparison of the occupancies of the intermediate R-loop state *I* for mismatches at positions 11 and 17 in the absence (upper panel) and presence (lower panel) of negative torque.

To explore the impact of seed mismatches in more detail, we plotted the measured R-loop formation times for selected torque values as a function of the mismatch position (Fig. 4E). Using a 10-fold increased R-loop formation time compared to the WT target as hypothetical threshold for a seed mismatch, we observed that the length of the seed region decreased with increasing torque from ∼6 bp at -6.7 pN nm to ∼15 bp at -3.4 pN nm. Predicting the R-loop formation time in absence of an external torque suggested that in this case the seed region would cover almost the entire target sequence (see black line in Figure 4E). To verify the prediction, we set up a fluorescence bulk solution assay based on a donor-quencher pair at the PAM distal end (Fig. 4F) and measured the R-loop formation kinetics in the absence of supercoiling for different mismatch positions (Figure 4G). The extracted mean times of R-loop formation confirmed the theoretical prediction (Figure 4E) demonstrating that the seed region in Cascade is mainly a product of the applied supercoiling, i.e. the bias of the free energy landscape. This can be intuitively understood by considering the different occupancies of the intermediate R-loop state *I* for PAM-proximal and distal mismatches, which determines the full R-loop formation rate. In absence of a bias, all R-loop lengths up to the mismatch are energetically equal and thus equally populated, which supports similar R-loop formation times (Figure 4H upper panel). In presence of a negative bias, the *I* state is energetically higher with respect to the *U* state and thus less populated for a PAM-proximal compared to a PAM-distal mismatch (Figure 4H lower panel). This provides a comparably slower transition to the *F* state for the PAM-proximal mismatch.

### Intermediate R-loop dynamics in presence of two mismatches

After verifying the random walk model for targets with single mismatches, we next tested whether it can be directly applied to describe the R-loop dynamics in presence of two mismatches. We produced double point mutants with the first mismatch located at positions 11, 13 or 14 and the second at position 17 within the locking deficient target (PAM-distal mismatches at positions 27-32). In this case R-loops could fluctuate between four possible states: *U*, the intermediate states before each mismatch *I* and *I*^*\**^ as well as *F*^*\**^ (Figure 5A). For the 11-17 double mismatch target, the *I* and the *I*^*\**^ states could be distinguished, but the transitions between the states were too fast to be resolved for the closer mismatch spacings (Figures 5B and 5D). State occupancies and transition dynamics were again torque-dependent (Figure 5C). Using the best-fit parameters from the single mismatch experiments, the random walk model predicted the measured state occupancies remarkably well (Figures 5B and 5C). A 4-state approximation of the recorded trajectories for the 11-17 double mismatch substrate (Figure 5D) allowed to extract the six transition rates between subsequent states. We obtained agreement of the extracted rates with the model predictions and the results of Brownian dynamics simulations (Figures 5E, S3D and S5) demonstrating that the random walk model could describe the R-loop dynamics also for multiple mismatches. Interestingly, the *F*^*\**^ state was less frequently visited as closer the two mismatches were positioned (Figure 5B) suggesting that the mismatch proximity influences the formation of the full R-loop.

**Figure 5.**
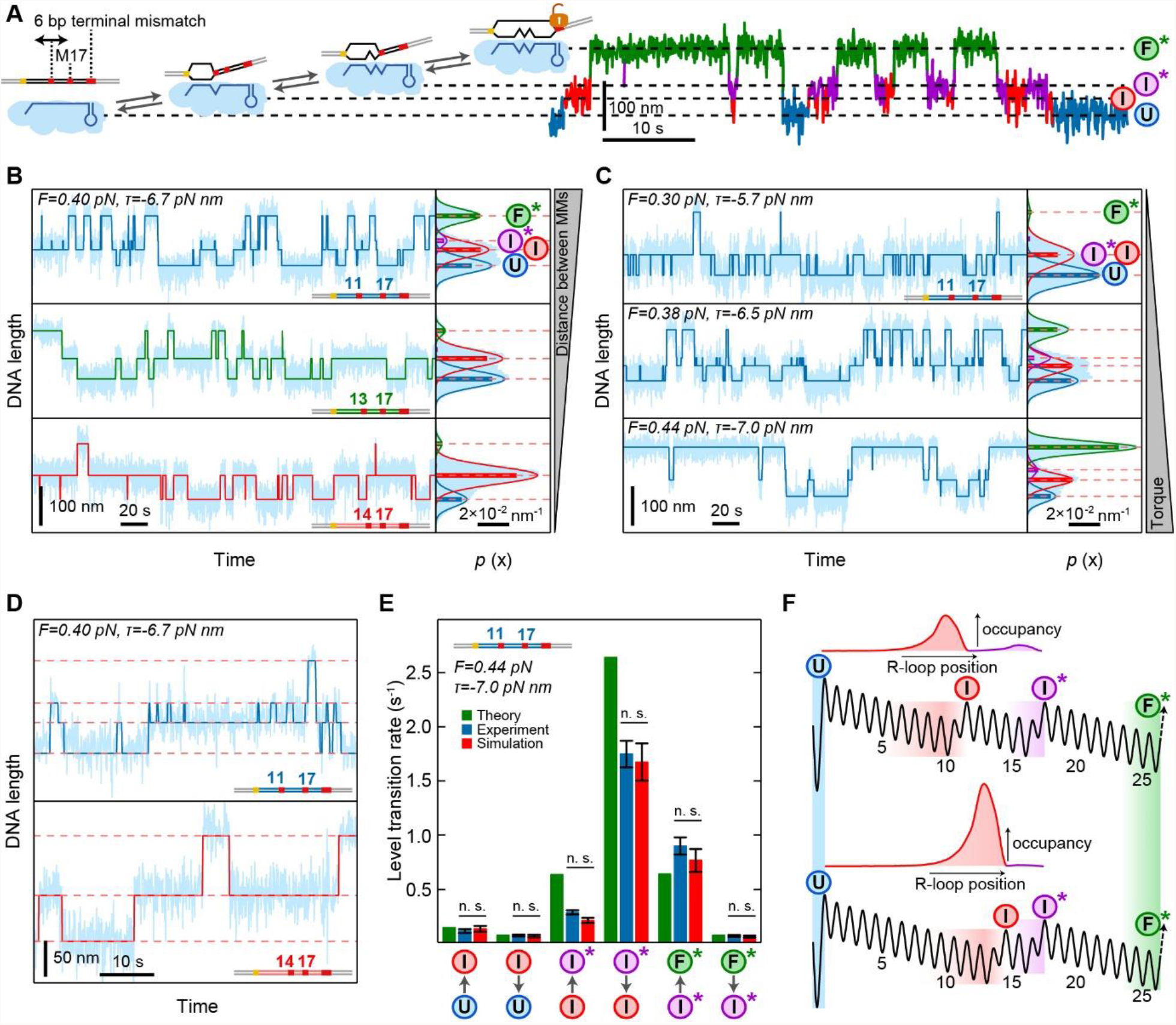
R-loop dynamics on locking-deficient targets containing two internal mismatches. (A) Sketch of the DNA construct with mismatched bases shown in red and the possible R-loop states: the unbound state (*U*), two intermediate R-loops (*I* before the first and *I*^*\**^ before the second mismatch) and the almost full but unlocked R-loop (*F*^*\**^). Sequential transitions between these states can be described by 6 different rates. Example trajectory of the transitions between the different states measured for a C:C mismatches. (B) Trajectories and histograms of the DNA length recorded for targets containing C:C mismatches with varying proximities at a torque of -6.7 pN nm (light blue). 4-state and 3-state (in case *I* and *I*^*\**^ states were indistinguishable) approximations of the trajectories in B and C are shown as dark lines. Solid lines in the histograms represent Gaussian fits experimental histograms, while horizontal dashed lines indicate the average DNA length of each state. Bars represent theoretically predicted occupancies using the parameters shown in SI Table 1. (C) Trajectories and histograms of the DNA length recorded for different applied torques using a target with C:C mismatch at positions 11 and 17 (light blue). (D) Enlarged view into the trajectories of the 11-17 and the 14-17 double-mismatch targets from (C) together with corresponding 4-state and 3-state approximations, respectively. (E) Theoretical (green), experimental (blue) and simulation-based (red) transition rates between adjacent states for the 11-17 double-mismatch target at a torque of -7.0 pN nm. (F) Model energy landscapes for R-loop formation for the 11-17 (top) and the 14-17 (bottom) double mismatch targets together with the normalized occupancies of the *I* and *I*^*\**^ states obtained from the model. Error bars in all subplots correspond to SEM.

### Proximity between double mismatches strongly influences R-loop formation

To investigate the influence of the proximity between two mismatches in detail, we studied locked R-loop formation on double C:C mismatch targets without terminal mismatches (Figure 6A). Predictions by the random walk model showed that a strong inhibition is obtained when combining two PAM-proximal mismatches (brown areas in Figure 6B, lower left corner) while a weak inhibition is obtained when combining two PAM-distal mismatches (blue areas in Figure 6B). However, a considerable inhibition is also obtained when combining two PAM-distal mismatches that are in close proximity (green tail along the diagonal in Figure 6B). This influence of the proximity between mismatches was only obtained for the random walk model but not for a simple addition of apparent free energy penalties (Figure 6B, lower upper right, Figures S6C and S6D) as applied in heuristic scoring schemes (Doench et al., 2016).

**Figure 6.**
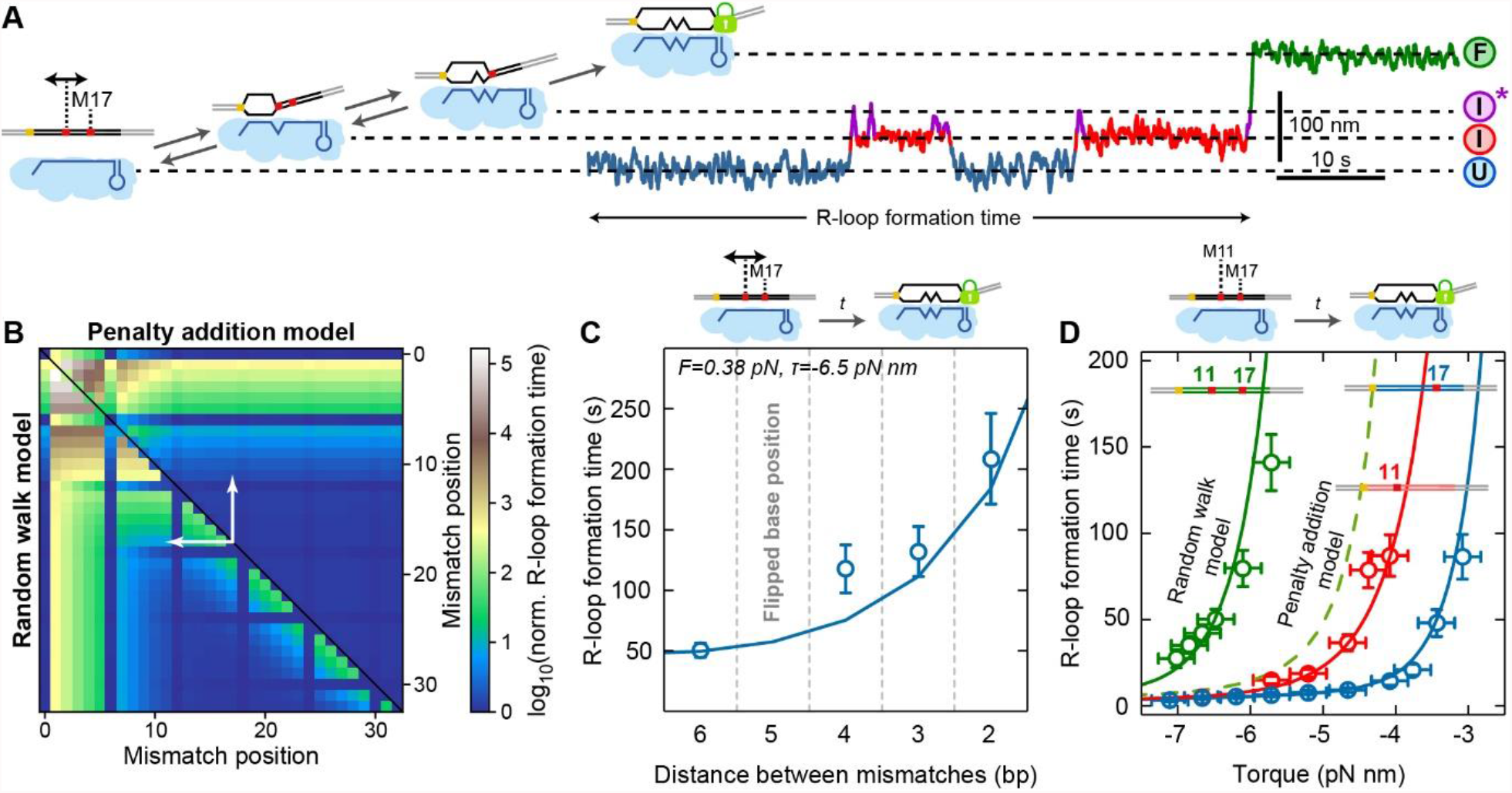
Locked R-loop formation on targets containing double mismatches. (A) Sketch of the DNA construct containing a first C:C mismatch at positions varying from 11 to 15 and a second C:C mismatch at position 17 (left). Example trajectory of full R-loop formation with the first mismatch located at position 11 containing two intermediate states *I* and *I*^*\**^. (B) Predicted R-loop formation time for double mismatches relative to the matching target using the penalty for a C:C mismatch and a torque of -6.5 pN nm. Either the penalty addition model (upper right half) or the random walk model (lower left half) was applied. For mutations at positions 6, 12, 18, 24 and 30 no penalties were considered due to the disrupted base-pairing in the crRNA-DNA hybrid. White arrows indicate the range of considered mismatch distances between positions 11 and 17. (C) Experimental R-loop formation times on targets with double mismatches with a variable position of the first mismatch and a fixed position of 17 for the second mismatch. The solid line represents the model prediction using the parameters from the single mismatch experiments (see SI Table 1). Error bars correspond to SEM. (D) R-loop formation time as function of torque for targets with single mismatches at positions 11 and 17 and the target containing both mismatches. Solid lines represent the model prediction using the parameters from the single mismatch experiments (see SI Table 1). The dashed line represents the prediction of the penalty addition model for the target with both mismatches. Error bars correspond to SEM; torque errors correspond to 0.25 pN nm.

To confirm the prediction, we next determined R-loop formation times as function of torque for a second mismatch at the fixed position 17 and distances to the first mismatch ranging from 2 to 6 bp (Figures S6A and S6B). The mean R-loop formation time at a given torque increased strongly with decreasing distance between the two mismatches (Figure 6C). This influence of the mismatch proximity was quantitatively predicted by the random-walk model using the parameters from single-mismatch experiments. Also, the torque dependence in case of two mismatches was quantitatively predicted by the model (Figure 6D). Altogether, the random walk model could successfully describe the non-trivial dependence of the R-loop formation kinetics on the mismatch distribution. Intuitively, this can be understood by considering that for close mismatches the *I* state is energetically lower than for distant mismatches, such that the occupancy of the *I*^*\**^ is lowered at the expense of the *I* state (Figure 5F). This provides in turn less frequent transitions to the *F* state for proximal compared to distal mismatches.

## DISCUSSION

In this study, we presented a highly comprehensive investigation of the R-loop dynamics within a CRISPR-Cas type I Cascade effector complex. Using direct measurements of the DNA untwisting during R-loop formation, we could uniquely resolve multiple R-loop intermediates on DNA targets with mismatches and carefully quantify their behavior as function of the positions and types of the mismatches.

Importantly, for the obtained data sets a unified biophysical model based on a random walk on a simplified one-dimensional energy landscape could be established. Mismatches were found to impose free energy barriers for R-loop expansion while applied supercoiling caused a global bias. Since the applied random-walk process was reversible, we could directly conclude that Cascade uses a kinetic target recognition mechanism governed by the kinetic competition between mismatch bypass and R-loop collapse. Increasing free energy barriers from mismatches disfavored R-loop progression and promoted its collapse (Figures 4A), which thus led to slower recognition kinetics of mismatched targets (Figure 4D). Such a kinetic target recognition process that is terminated by an irreversible locking step is fundamentally different from typical affinity-based mechanisms (Bisaria et al., 2017). In principle, given sufficient time, any target can become recognized. This sets important constraints when discussing specificities of effector complex variants (Kleinstiver et al., 2016; Lee et al., 2018; Slaymaker et al., 2016; Wu et al., 2018).

The success of the random walk model implies that R-loop formation samples the continuum of available R-loop lengths and that unlocked R-loop intermediates dynamically extend over a range of different lengths (Figure 2F). This is in contrast to a recent report identifying only 2-discrete states during R-loop formation by Cas9 (Ivanov et al., 2020). Furthermore, our data is in agreement with a very simple form of the energy landscape for Cascade that is in absence of mismatches essentially flat (Figure 1B). Except for a small positive intrinsic bias, external supercoiling sets typically the overall bias of the landscape thus driving R-loop formation. Our modelling also provided a first insight into the time scale of the single base-pair R-loops steps that were found to be in the millisecond to sub-millisecond range (Tables S1 and S2).

An important finding of our data and our modelling is that the seed sequence observed for all R-loop forming CRISPR-Cas effector complexes (Künne et al., 2014) is at least partially a biophysical consequence of the external DNA supercoiling. At negligible bias of the energy landscape, mismatches had a rather position-independent impact such that an extended seed was observed for Cascade targeting (Figure 4E). In contrast, at sufficient negative bias of the energy landscape induced by supercoiling, the target recognition was only little affected by PAM-distal mismatches which strongly limited the seed extension. In this case, R-loops that were stopped by the PAM-distal mismatches reached sufficiently low free energy values such that they had only low chances to collapse.

The observed torque-dependence at *in vivo* relevant torque values (Figure 4D) can explain observations of a short well-defined seed region *in vivo* (Fineran et al., 2014; Semenova et al., 2011) in contrast to more relaxed seed conditions *in vitro* in absence of supercoiling (Jung et al., 2017; Rutkauskas et al., 2015). The absence of a well-defined seed is in agreement with structures of Type I-E Cascade complexes (Hayes et al., 2016; Mulepati et al., 2014; Xiao et al., 2017) where specialized seed motives were not observed. Generally, seed regions for RNA-guided nucleic acid recognition can also have structural determinants, as e.g. observed for Cas9 (Jiang et al., 2016). In the view of our model, this would be achieved by specific interactions of the R-loop with the protein complex that would establish an intrinsic bias of the free energy landscape already in absence of supercoiling.

The supercoil- and position-dependence of the mismatch impact directly affects the specificity of the target recognition process. CRISPR-Cas effector complexes have predominantly evolved for activity on negatively supercoiled DNA as typically found in prokaryotic cells (Bauer, 1978; Dorman, 2019). These conditions would provide less stringent specificities but accelerated target recognition kinetics. In genome engineering applications of eukaryotic cells, however, only moderate supercoiling levels are present, which would make the target recognition more specific but also slower. Noticeably, the supercoiling in eukaryotic cells is highly locus specific (Corless and Gilbert, 2017; Kouzine et al., 2013; Naughton et al., 2013) which should be considered when developing improved locus-specific off-target predictors.

From the R-loop dynamics on targets with single mismatches we could directly obtain free energy values for individual mismatch penalties. The position independence of the penalties suggested that our random-walk model accounted correctly for the position-dependent bias observed previously (Jung et al., 2017). The found free energy penalties for the three different mismatches were similar to apparent penalties determined in high-throughput *in vitro* binding experiments (Jung et al., 2017) but different to mismatch penalties within DNA duplexes (Table S1, (SantaLucia and Hicks, 2004)). Most importantly, we could directly apply the single mismatch penalties to quantitatively predict the R-loop formation dynamics on targets with two mismatches. Particularly, the model could correctly describe an increased inhibition of R-loop formation with decreasing mismatch distance (Figure 6C). This proximity-dependence can be intuitively understood by considering the corresponding energy diagrams (Figure 5F). Alternatively, one can consider the region between the two mismatches as seed region for full R-loop formation when being in the *I* state. An easier escape from the seed region (*I*^*\**^ state) back to the *I* state is then possible for proximal compared to distal mismatches (see also Figure 4H).

Given its applicability to double mismatches, we expect that our model can be easily extended to larger numbers and any types of mismatches. This way an off-target predictor based on biophysical modelling would be obtained for which our careful modelling of the R-loop dynamics provides solid validation. Missing parameters for such a predictor would be the penalties of all possible mismatches corresponding to at least 48 parameters due to 12 different mismatch types and 4 different nearest neighbor base pairs onto which a mismatch can stack on each side (SantaLucia and Hicks, 2004). Stringent analysis of high-throughput binding data on *in vitro* target libraries (Boyle et al., 2017; Jung et al., 2017) with our random walk model should allow for such a comprehensive parametrization. Overall, we expect that the resulting biophysical model will considerably better and most importantly rationally integrate the types and locations of different mismatches as well as proximity-based effects than current off-target predictors. As a result it should enable a more rational selection of target sequences for a precise programming of DNA editing tools with low off-target probabilities. After determination of the corresponding energy landscapes, our model should be readily applicable to all other R-loop forming CRISPR-Cas effector complexes such as Cas9 and Cas12a, due to the high mechanistic similarities in target recognition. Experiments similar to the ones presented here performed on other CRISPR-Cas nucleases will help to determine the impact of structural peculiarities and R-loop-protein interactions on the intrinsic energy landscapes.

## METHODS

### One-dimensional random-walk model for R-loop formation

The R-loop dynamics was modeled as a random walk on a one-dimensional 1 bp lattice within a simplified free energy landscape based on the energy parameters *ΔG*_*ini*_, *ΔG*_*bias*_, *ΔG*_*lock*_ and *ΔG*_*MM*_ as described in the main text (see also Figure 1B). The rate model of the random walk was parameterized based on the principle of detailed balance which relates the ratio of the forward (indicated by ‘+’) and the backward (indicated by ‘-’) rate constants between subsequent positions *n* and *n* + 1 to the free-energy difference *G*_*n*+1_ − *G*_*n*_ between these positions:

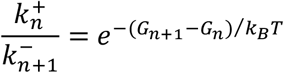

For the unbiased energy landscape with *ΔG*_*bias*_ = 0 (Figure 1B, top), we assume an equal rate *k*_*step*_ for all transitions between R-loop states (i.e. equal kinetic barriers) except for transitions from the *U* (and *F*) states. This excludes any sequence dependence of the stepping rates. PAM binding is comprised in the formation step of the first R-loop base pair without a distinct kinetic barrier for dissociation from the PAM. Based on these considerations the rate for R-loop initiation is given as:

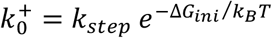

The dependence of R-loop initiation on the Cascade concentration *c* was included by considering the contribution of the chemical potential of the Cascade complexes to *ΔG*_*ini*_:

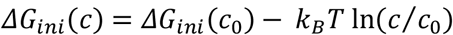

where *ΔG*_*ini*_(*c*_0_) is the standard initiation penalty at a reference concentration *c*_0_.

For a mismatch between crRNA and DNA target strand at position *m*, we assume that the rate limiting step for mismatch establishment is the disruption of the DNA base pair, such that it occurs at the normal R-loop extension rate 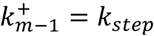. Detailed balance provides in this case an increased rate for R-loop retraction that eliminates the mismatch:

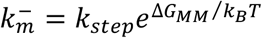

in agreement with the rate-limiting step being facilitated by destabilized base-pairing in the heteroduplex as also indicated by the lowered kinetic barrier at the mismatch position in Fig. 1B.

The applied negative supercoiling causes a constant bias of the energy landscape per bp in the regime where the DNA length decreases linearly with the applied turns. The torque *τ*, which is the quantitative parameter of how the applied supercoiling stresses the DNA helix, is set by the applied force in the magnetic tweezers experiments. It was estimated as described before (Maffeo et al., 2010). The bias *ΔG*_*bias*_ per bp from the torque equals the work done against the torque:

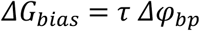

where *Δφ*_*bp*_ = 0.515 rad(≙ 29.5°) is the angle by which the DNA becomes untwisted per R-loop expansion by 1 bp (Szczelkun et al., 2014). Assuming that the transition barrier is centered between two subsequent R-loop positions, R-loop expansion and retraction would both be affected by half of the bias providing the following corrections of all forward and backward rates for the acting torque:

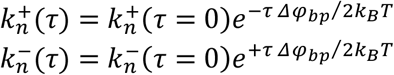

with *n* being any valid position of the energy landscape. These definitions provide a full parameterization of the rate model that describes the random walk. Mean transition times between any starting state and any end state were calculated by solving the first passage problem for this model (see Supplementary Note 1). To this end, transmissive boundaries were placed at the positions of end states and a single particle was added to the system. Upon arrival of the particle at a transmissive boundary it was instantaneously set to the start state. The mean transition time was then calculated from the steady-state flux of the single particle inside the rate landscape (see Supplementary Note 1). For intermediate R-loop states that are dynamic and extend over several base pairs, the position with the lowest free energy was taken as the state position. For DNA targets with a continuous stretch of mismatches at the PAM-distal end, the energy landscape was cut off at the first mismatch position corresponding to an infinite free energy at this position.

For kinetic random walk simulations of R-loop length fluctuations (see Figure S3), we constructed the energy landscape for a given target and calculated for each step of the lattice the forward and backward stepping rate constants 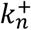 and 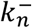. Per simulation time step *Δt*, a single bp forward or backward step was taken with probability 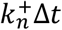 or 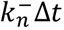, respectively. *Δt* was chosen sufficiently small, such that the stepping probabilities were much smaller than one.

### Estimation of the DNA torque in *E*.*coli* cells

Typical superhelical densities *σ* (i.e. the number of added superhelical turns per helical turns of the relaxed DNA) found in *E*.*coli* cells range from −0.06 to −0.029 (Brouns et al., 2018; Higgins and Vologodskii, 2015). For plasmid DNA added superhelical turns are partitioned between writhe and twist at a ratio of 0.8 to 0.2 (Ubbink and Odijk, 1999). Thus, the superhelical density contributing to the DNA twist is:

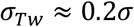

The torque *τ* in a twisted semiflexible polymer of length *L* can be calculated from:

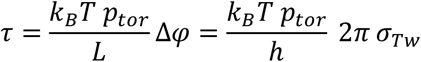

where *p*_*tor*_ is the torsional persistence length and *Δφ* the twist angle. The number of helical turns within *L* is given by *N*_*helix*_ = *L*/*h*, where *h* = 3.5 nm is the helical pitch of B-form DNA. The twist angle is then given by *Δφ* = 2*π σ*_*Tw*_*N*_*helix*_. Inserting these relationships in the torque equation gives:

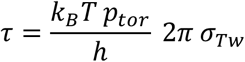

Using *p*_*tor*_ = 100 nm (Bouchiat et al., 1999; Kauert et al., 2011; Lipfert et al., 2010), we get for the typical superhelical densities in *E*.*coli* cells torques *τ* between −8.9 and −4.3 pN nm.

### DNA and proteins

The 2.1 kbp DNA constructs with additional biotin- and digoxigenin modified attachment handles at either end used in the magnetic tweezers experiments were prepared as previously described (Luzzietti et al., 2012; Szczelkun et al., 2014). For each DNA target presented in this study a 73 bp blunt ended oligonucleotide duplex carrying the 35 bp long target sequence was cloned into the SmaI site of a pUC19 plasmid. From the plasmids, 2.1 kbp DNA fragments containing the targets as well as SpeI and NotI restriction enzyme sites at either end were produced by PCR (primers 1 and 2 in Table S3). Biotinylated and digoxigenin-modified ∼1.2 kbp DNA fragments were produced by PCR from pBluescript II SK+ with its multiple cloning site located approximately in the center of the fragments (primers 3 and 4 in Table S3). The biotinylated and digoxigenin-modified fragments were digested with SpeI and NotI, respectively to yield modified ∼600 bp attachment handles. Following digestion of the 2.1 kbp target fragment with both restriction enzymes, it was ligated with the handles to form the final 3.3 kbp DNA construct used in magnetic tweezer experiments. For the production of complexes with different crRNAs all spacer variants were introduced into the produced vector pACYCminCR-Eco31I/SapI through the SapI and Eco31I sites using synthetic oligonucleotides with corresponding single stranded overhangs (Table S4) (Songailiene et al., 2019). Cascade complexes with different crRNAs were expressed in *E*.*coli* BL21 (DE3) cells and purified as described (Sinkunas et al., 2013) using the pACYC-minCR derivatives (CmR) instead of pACYC plasmid with homogeneous CRISPR region pCRh.

### Magnetic tweezers experiments

R-loop-formation experiments were performed in 20 mM Tris-HCl (pH 8.0), 150 mM NaCl and 0.1 mg ml^-1^ BSA at 170 pM (for experiments described in figures 2 and 3) or 0.5 nM (for experiments described in figures 4, 5 and 6) Cascade using a custom-built magnetic tweezers setup (Klaue and Seidel, 2009) at room temperature (25 °C). DNA molecules were bound to 0.5 μm streptavidin-coated magnetic beads (MasterBeads; Ademtech) and added into the antidigoxigenin-coated flow cell to form tethers at the bottom surface (Luzzietti et al., 2012; Schwarz et al., 2013). Single supercoilable molecules were selected (Rutkauskas et al., 2017). The DNA length was determined at 120 Hz by videomicroscopy and real-time GPU-accelerated three-dimensional particle tracking (Huhle et al., 2015) from the position of the magnetic bead with respect to a surface-bound non-magnetic reference bead (Dynospheres; Invitrogen). Forces were calibrated from the lateral fluctuations of the DNA-tethered magnetic beads (Daldrop et al., 2015). Torque values were calculated based on previous theoretical work (Maffeo et al., 2010; Schöpflin et al., 2012). For experiments in which dynamic sampling of R-loop intermediates in absence of locking was investigated (Figures 2, 3 and 5), Cascade was added to the flow cell and the DNA molecule was negatively supercoiled (see Figure S1A). The number of negative turns depended on the applied force and on the change of supercoiling following the formation of the R-loop. Generally we aimed that DNA length transitions occur around half the relaxed molecule length at the given force and thus in the linear regime of the supercoiling curve (Rutkauskas et al., 2015, 2017). This way the torque on the DNA stayed approximately constant. For experiments where R-loops became locked (Figures 4 and 6), R-loop formation was induced as described before. To remove Cascade complexes with locked R-loops, the DNA molecule was positively supercoiled to (+10-14 turns, depending on the force and on the length of the individual DNA molecule) and the force was increased to ∼2-3 pN to provide a high positive torque that would ‘wring out’ the R-loop. R-loop dissociation was seen as a sudden DNA length increase (see Figures S1B-S1E for the full procedure). R-loop formation-dissociation cycles were constantly repeated to obtain ≥25 individual events per applied condition.

### Fluorescence bulk solution experiments

All oligonucleotides for the zero torque measurements are shown in Table S5 and were purchased HPLC-purified from Sigma-Aldrich. Shipping concentrations of 100 μM were evaluated by measuring the absorbance at 260 nm using a P-330 NanoPhotometer (Implen). Complementary strands were then annealed at a concentration of 1 μM in buffer containing 10 mM Tris−HCl (pH of 8.0), 50 mM NaCl, and 1 mM EDTA and slow cooling from 95 to the storage temperature of 4 °C at 1 K/min.

All measurements were performed in a temperature controlled Cary Eclipse at 25 °C in 1500 μL cuvettes. Before each measurement, cuvettes were rinsed 5 times with ethanol, 5 times with mili-Q, incubated overnight in 2% Hellmanex 3 solution, and again rinsed 5 times with ethanol and 5 times with mili-Q.

In the beginning, a 1350 μL solution containing the double stranded DNA was measured for 600 s to obtain the ground level (9/10 of mean signal amplitude). Afterwards the reaction was started by quickly adding 150 μL of solution containing Cascade. Reaction conditions were 10 nM of dsDNA and 2 nM Cascade in a buffer containing 20 mM Tris−HCl (pH of 8.0), 150 mM NaCl and 0.1 μg/μL BSA.

The negative control (no protein added) was then subtracted from the measured trajectories. The fluorescence signal was then fitted to a sum of three exponentials of the form:

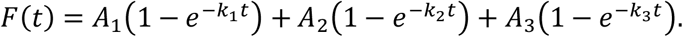

The quantity of interest is then; the mean time it takes to overcome all three steps:

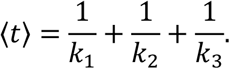

### Data analysis

DNA length trajectories recorded at 120 Hz were smoothed with a sliding average to 7.5 Hz. Transitions between the different R-loop states to generate 2-, 3- and 4-state approximations of the R-loop trajectories were obtained by hidden Markov modeling using the vbFRET software package (Bronson et al., 2009). From the discrete-state trajectories, dwell time distributions and transition rates were extracted using MATLAB (MathWorks) and LabVIEW (National Instruments). This included the generation of cumulative dwell time distributions for individual states, which were fitted to single exponential functions to obtain mean dwell times and the corresponding transition rates. For the latter, the transition probabilities to neighboring states were correspondingly considered. For experiments in which dynamic sampling of R-loop intermediates was investigated (Figures 2, 3 and 5), trajectories of at least 3000 s were recorded for each condition including typically ∼1000 transitions. For experiments where R-loops became locked (Figures 4 and 6), ≥25 locking events were obtained for each condition to determine mean R-loop formation times. All rate and time error bars represent the standard error of the mean. Particularly, the error of mean dwell times was calculated by dividing the mean time by the square root of the number of events.

To verify that the temporal resolution of our nanomechanical system (bead on supercoiled DNA was sufficient to resolve the extracted R-loop transitions, we employed Brownian dynamics simulations to simulate the magnetic tweezers measurement of dynamic R-loop sampling. We first employed kinetic random-walk simulations to simulate trajectories of the R-loop dynamics (R-loop length over time) on a one-dimensional 1 bp lattice within the corresponding energy landscape (see model description above) using the experimentally obtained parameters (Table S1). Using the slope of the supercoiling curve at the corresponding force, we converted the R-loop length into an expected equilibrium extension *z*_*DNA*_(*t*) of the DNA. We furthermore modelled the diffusive fluctuations of the magnetic bead and its response to DNA extension changes by one-dimensional Brownian dynamics simulations (Daldrop et al., 2015). In brief, a deviation *Δz* of the fluctuating bead from the equilibrium extension, caused a back-driving drift force *F*_*drift*_ = −*κ Δz* = *z*_*bead*_ − *zD*_*NA*_(*t*) due to the stretch elasticity *κ* of supercoiled DNA (comprising components of the DNA twist elasticity and the entropic elasticity of stretched DNA). Per time increment of the simulation *Δt, F*_*drift*_ caused a displacement of *Δz*_*drift*_ = *Δt v* = *Δt F*_*drift*_/*γ*, with *v* being the steady state drift velocity of the bead inside the viscous medium for low Reynolds numbers. The viscous drag coefficient *γ* of a spherical particle with radius *R* inside a medium with viscosity *η* was given by the Stokes formula *γ* = 6*πηR*. Within *Δt*, we furthermore considered random diffusive displacements *Δz*_*diff*_ that were drawn from a Gaussian distribution with zero mean and a variance of ⟨*Δz*_*diff*_^2^⟩ = 2*D Δt* with *D* = *k*_*B*_*T*/*γ* being the diffusion coefficient for the particle. By successively updating the bead position *z*_*bead*_ by a total displacement of *Δz*_*diff*_ + *Δz*_*drift*_ per time increment, we obtained the magnetic bead fluctuations in response to the R-loop length fluctuations (see light red trajectory in Figure S3A-S3D and Figure S5). The spring constant *κ*∼0.01 pN/nm describing the stretch elasticity of supercoiled DNA as well as the effective hydrodynamic bead radius *R*∼800 nm were obtained from power spectral density analysis of corresponding experimental trajectories of the length fluctuations of supercoiled DNA (Daldrop et al., 2015).

We next identified for the simulated magnetic tweezers trajectories transitions between different R-loop intermediates using vbFRET and compared them with the actual transitions of the R-loop length simulations. We could correctly identify transitions for simple R-loop intermediates (as in Figure 2, main text) as well as for the 3-state transitions observed for sufficiently strong single mismatches (C:C and C:T, see Figure 3, main text, and Figure S3A and S3B). For targets containing a single C:A mismatch we observed a considerably slower collapse of the full R-loop state *F*^*\**^ (rate *k*_4_) compared to the C:T and C:C mismatches (Figures 3B and S3E) despite the anticipated independence of the R-loop collapse on the mismatch strength. We therefore hypothesized for this weak mismatch that transitions over the mismatch between the *I* to the *F*^*\**^ states (rates *k*_3_, *k*_4_) were too fast to be reliably detected given the temporal resolution of our setup. To correct for the undetected transitions between *I* and *F*^*\**^ states we increased the R-loop collapse rate *k*_4_ to the level measured for C:C and C:T mismatches (0.8 − 1.2 *s*^−1^), which proportionally also increases the R-loop formation rate *k*_3_ and used the adjusted rates for characterizing the mismatch penalty (see Figure S3E). We additionally carried out simulations using the adjusted rates. Simulations of the R-loop length fluctuations provided the expected fast transitions (see Figure S3E, green data points). The simulated magnetic tweezers trajectories provided significantly lower transition rates that agreed with the experimental rates, supporting a correct adjustment of the rates for this weak mismatch. The correction procedure was also applied for extracting transition rates for the dynamic sampling of R-loop intermediates in case of double mismatches (positions 11, 17, Figure 5E and Figure S5). While transitions between *U* and *I* as well as between *I*^*\**^ and *F*^*\**^ were correctly reproduced by the simulations, part of the fast transitions between the *I* and *I*^*\**^ states were not detected (Figure S5). Transition rate comparisons obtained from each trajectory are represented in Figure 5E.

## SUPPLEMENTAL FIGURES

**Figure S1.**
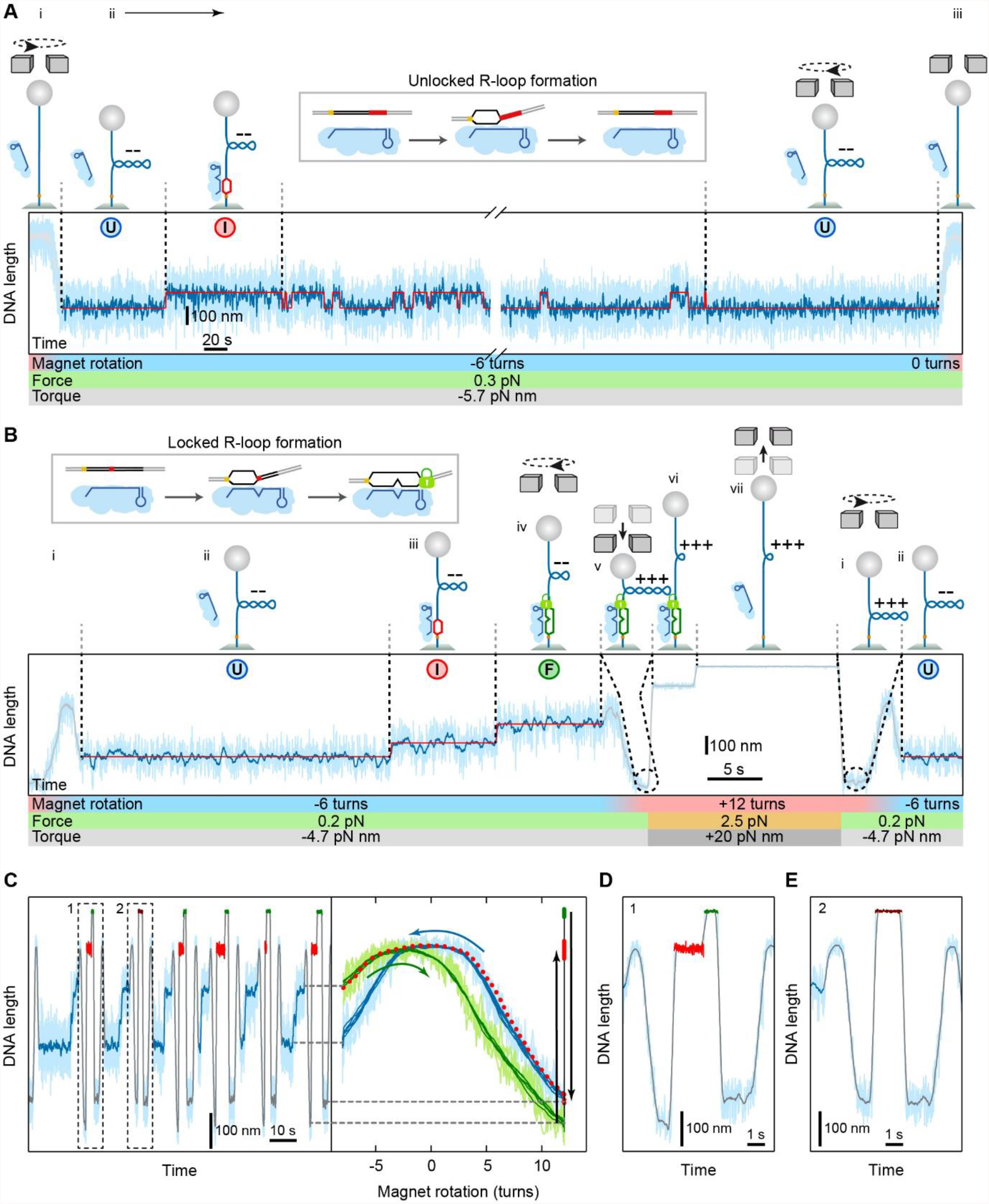
Detailed representation of DNA length, applied turns, force and torque during the different R-loop formation experiments using magnetic tweezers. Related to Figures 1-6. (A) Experiment to study the R-loop dynamics on a target that does not lock due to ≥6 bp PAM distal mutations (20 bp for the depicted example or 12 matching bp). The experiment starts with supercoiling the DNA molecule from 0 turns to -3 to -8 negative turns at constant magnet position (i). The molecule length reduces due to the DNA writhe. The magnetic field force is kept constant providing a constant negative torque acting on the DNA that is controlled by the force (ii). Thermally driven R-loop formation and collapse events seen as discrete changes of the DNA length (see two state approximation shown as red line as well as cartoons on top) are followed over a sufficiently long time. Finally, negative DNA supercoiling is removed by turning the magnets to the initial 0 turns leading to a DNA length increase (iii). The light blue trajectory represents the raw DNA length data collected at 120 Hz, the dark blue trajectory the DNA length after sliding-window filtering to 7.5 Hz. Red lines represent the two-state approximation of the trajectory. (B) Experiment to study the R-loop formation kinetics in case of locking. In order to measure multiple R-loop formation events Cascade needs to be dissociated from the locked state. First, R-loop formation is facilitated by applying negative turns at low force of 0.1 – 0.45 pN (transition from i to ii). After observing formation of *I* state (iii) and subsequent locked R-loop formation (seen as a sufficient DNA length increase corresponding to the *F* state iv), the R-loop dissociation is induced by supercoiling the DNA towards positive turns (v) followed by the application of a higher force of ∼2.5 pN (vi) that provides an increased positive torque. R-loop dissociation is observed as a discrete DNA length increase (vii). For a new R-loop formation experiment, the force is again lowered and the DNA is supercoiled to the initial negative turns (i and ii). (C) Example of multiple R-loop formation-dissociation cycles for a substrate containing a mismatch at position 7. Left panel represents the DNA length time trajectory depicted as in (B). Red areas of the trajectory represent R-loop at high force and positive supercoiling as vi in B. Green areas of the trajectory represent DNA length after the R-loop dissociation as vii in B. Right panel represents DNA length dependency on magnet rotation. Blue curve shows magnet rotation after R-loop dissociation from positive supercoiling to negative (i in B). After R-loop formation (seen as an abrupt jump of DNA length in left panel or transition from ii to iii to iv in B) DNA is being supercoiled from negative turns to positive turns and is shown as a green curve (transition from iv to v in B). In some rare cases the full R-loop was formed but remained unlocked (depicted as brown area in left panel). In this case the R-loop collapse occurred around zero turns where it could not be seen (red dashed curve in the right panel). These events were not considered in further data analysis. To verify whether the R-loop was locked, we monitored the expected sudden DNA length increase upon R-loop dissociation as well as the shift of the rotation curves at positive turns expected for stable locked R-loops. (D) and (E) enlarged views of the dissociation trajectories from C. In D R-loop presence is observed as lower magnetic bead position after DNA is positively supercoiled (v in B) and as a presence of two distinct states after the increase of force (red and green areas, vi and vii in B). In case of R-loop dissociation during transition through 0 turns (red dashed curve in right panel of C) magnetic bead does not go as low as in the presence of R-loop (i in B) and only one state is observed after the increase of force (brown area in left panel of C, vii in B) as represented in E.

**Figure S2.**
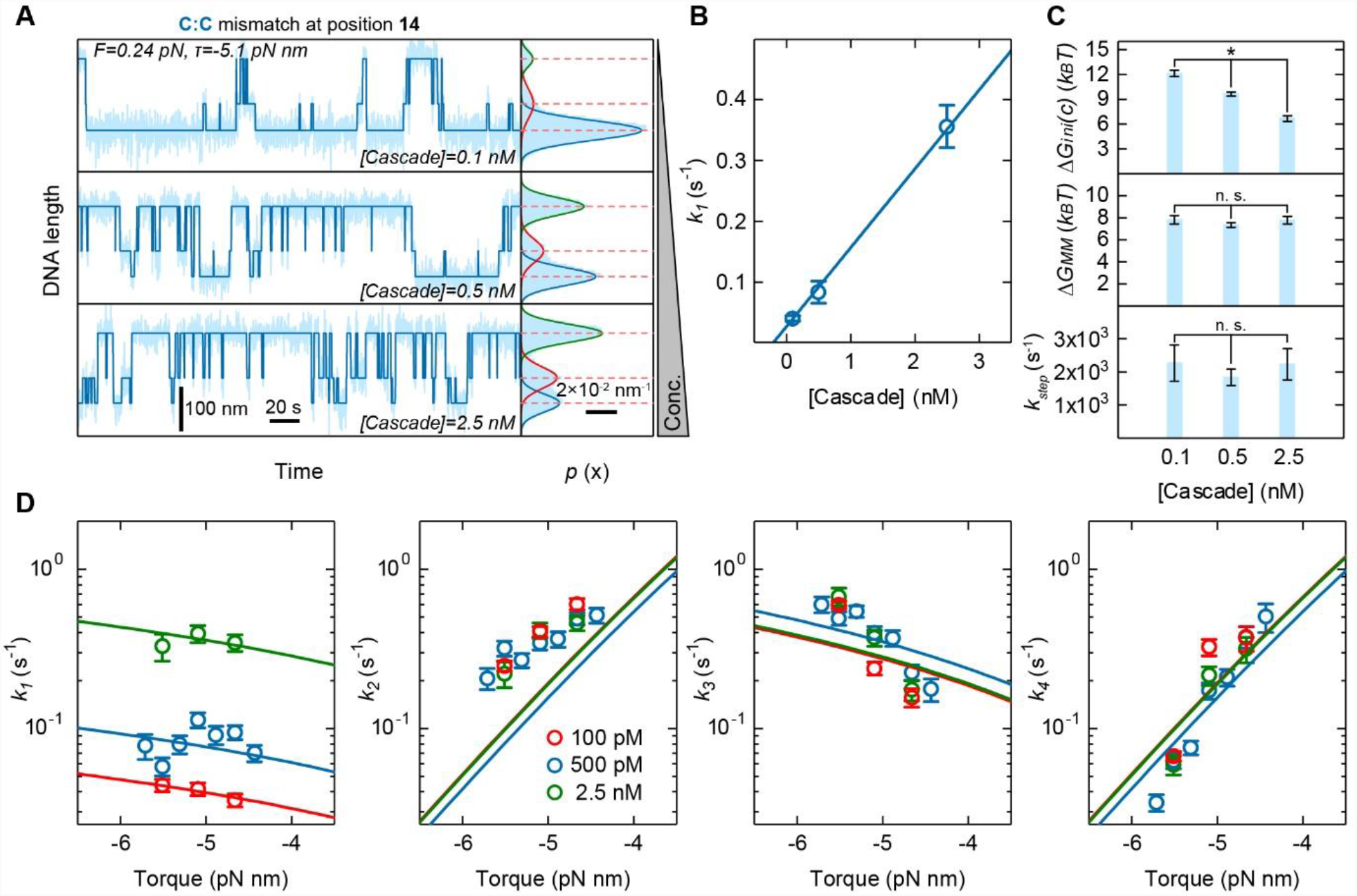
R-loop dynamics in presence of a single mismatch measured at different concentrations. Related to figure 3. (A) DNA length trajectories and occupancies measured at different Cascade concentrations on a target containing a C:C mismatch at position 14 and 6 PAM-distal mismatches. With increasing concentrations the full R-loop state *F*^*\**^ becomes increasingly populated. (B) Intermediate R-loop formation rate *k*_1_ as function of the Cascade concentration at -4.7 pN nm (open circles). The blue line represents a linear fit to the data. (C) Best fit parameters obtained for the different concentrations from the fits of the transition rates between the different R-loop intermediates (see D). (D) Experimental transition rates as function of torque for the different Cascade concentrations (open circles). Global fits to all rates at the given concentrations are shown as solid lines.

**Figure S3.**
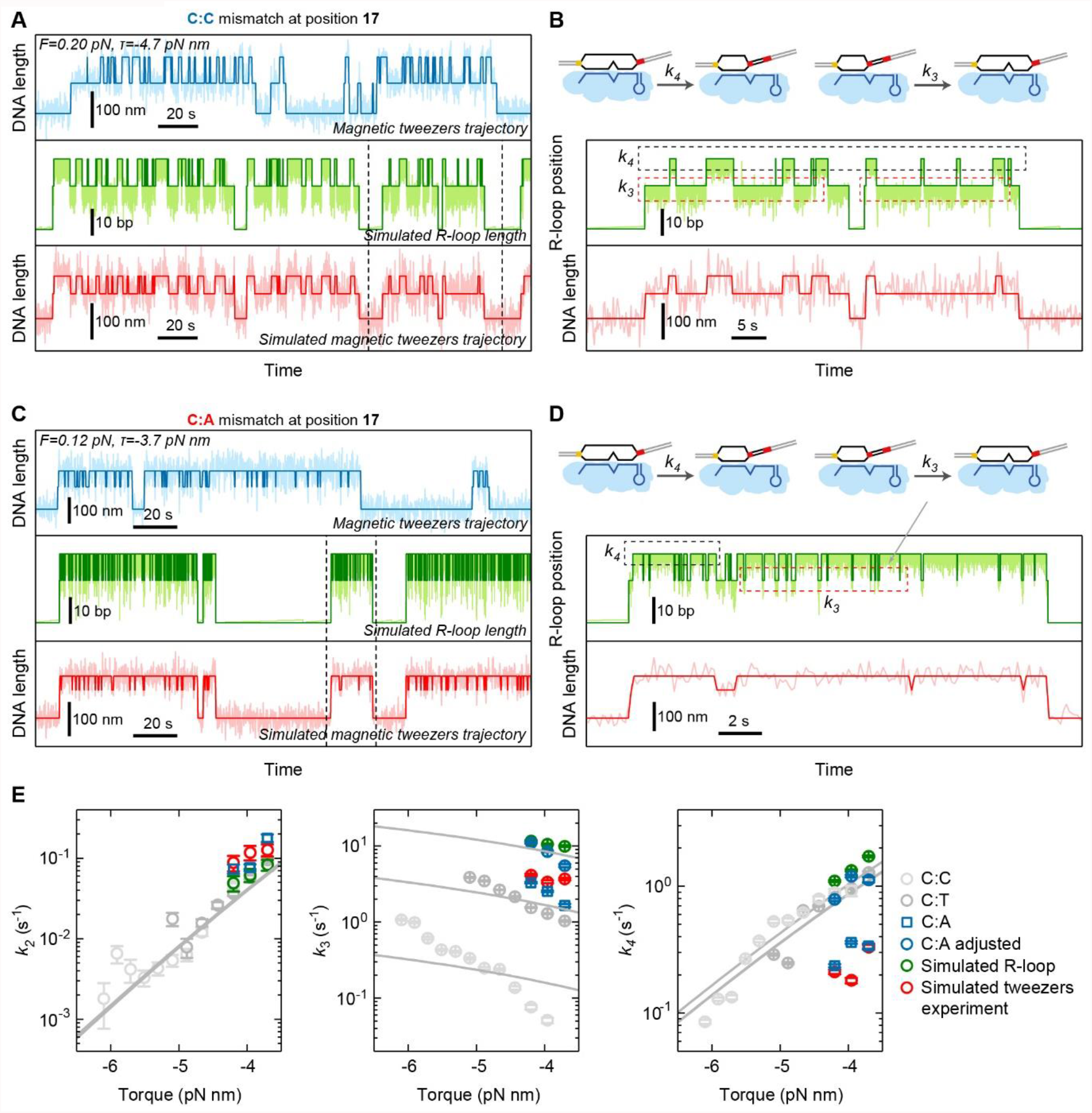
Simulations of R-loop length and magnetic tweezers experiments for single mismatch targets. Related to Figure 3. (A) and (C) Comparison between trajectories of a magnetic tweezers experiment (blue), a random walk simulation of the R-loop length (green) and a Brownian dynamics simulation of a magnetic tweezers experiment (red) for a target containing C:C mismatch (A) and C:A mismatch (C) at the position 17 and 6 terminal mismatches. Light colors show the raw results of the experiments and simulations while dark colors show corresponding 3-state approximations. (B) and (D) Enlarged view into the simulated trajectories from areas separated by dashed lines in (A) and (C) correspondingly depicting transitions extracted by the 3-state approximations of the simulated magnetic tweezers trajectories. Dashed boxes depict transitions contributing to *k*_3_ and *k*_4_. (E) Comparison of the extracted rates for measured and simulated trajectories. Measured *k*_3_ rate was adjusted by the ratio between measured *k*_4_ rates for C:C, C:T and C:A mismatches. Assuming *k*_4_ for C:A mismatch is in the same range as for C:C and C:T, C:A values were shifted upward. The same ratio was used to shift *k*_3_ values. After the adjustment, determined mismatch penalty values were used to perform Brownian dynamics simulations and to compare rates from the simulation experiment and magnetic tweezers experiment.

**Figure S4.**
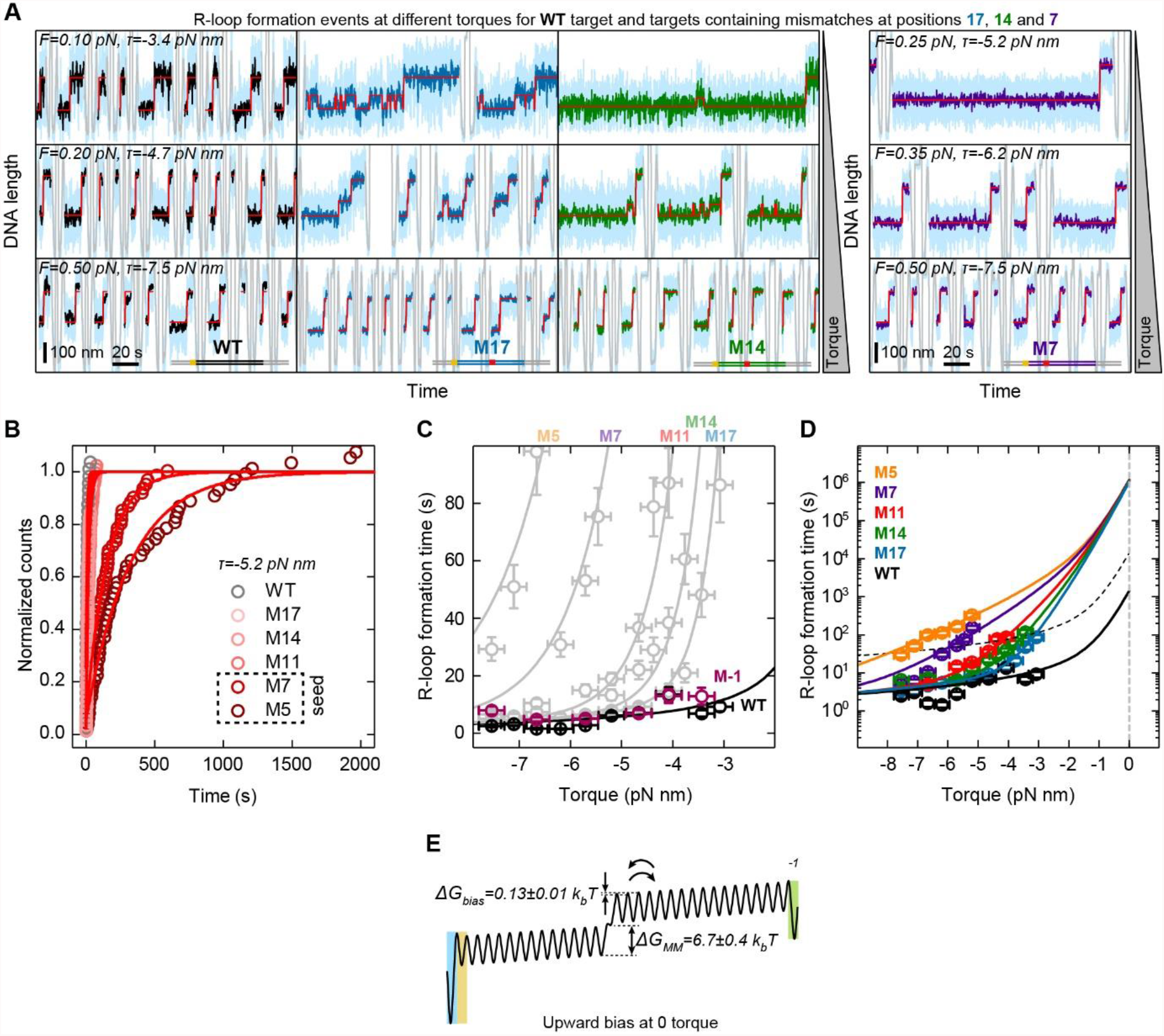
Additional data for locked R-loop formation on targets with single internal mismatches. Related to Figure 4. (A) Example trajectories for R-loop formation on targets containing a single C:C mismatch at various positions as indicated). Colored sections represent the actual R-loop formation events, while gray sections correspond to changes of the DNA supercoiling. The width of a formation event is roughly proportional to the R-loop formation time, such that a visual impression of the R-loop formation time for the different mismatches and torques can be obtained. Light blue trajectories represent raw magnetic tweezers data collected at 120 Hz, dark trajectories represent the data after sliding-window filtering to 7.5 Hz, red lines represent 2-state approximations of the trajectories for WT and M7 targets (the intermediate state is too short lived to be observed for M7) and 3-state approximations for M17 and M14 targets. (B) R-loop formation kinetics for the different targets at a torque of -5.2 pN nm (open circles) represented as normalized event count over time of the event occurrence. Single exponential fits to the data are shown as solid lines. (C) Comparison of the torque dependence of R-loop formation for the WT target and a target with a PAM mutation at position -1 (see Table S3). Data shown in gray is from Figure 4 (main text). (D) Torque dependence of the R-loop formation times for the different targets with single internal mismatches plotted with a semi-logarithmic time scale including an extrapolation of the fit curves to zero torque. Noticeable, the difference between the different targets with single mismatches vanishes at zero torque, while a strong difference to the WT target persists. (E) Depiction of the positively biased energy landscape of the R-loop formation for the target with the mismatch at the position 15.

**Figure S5.**
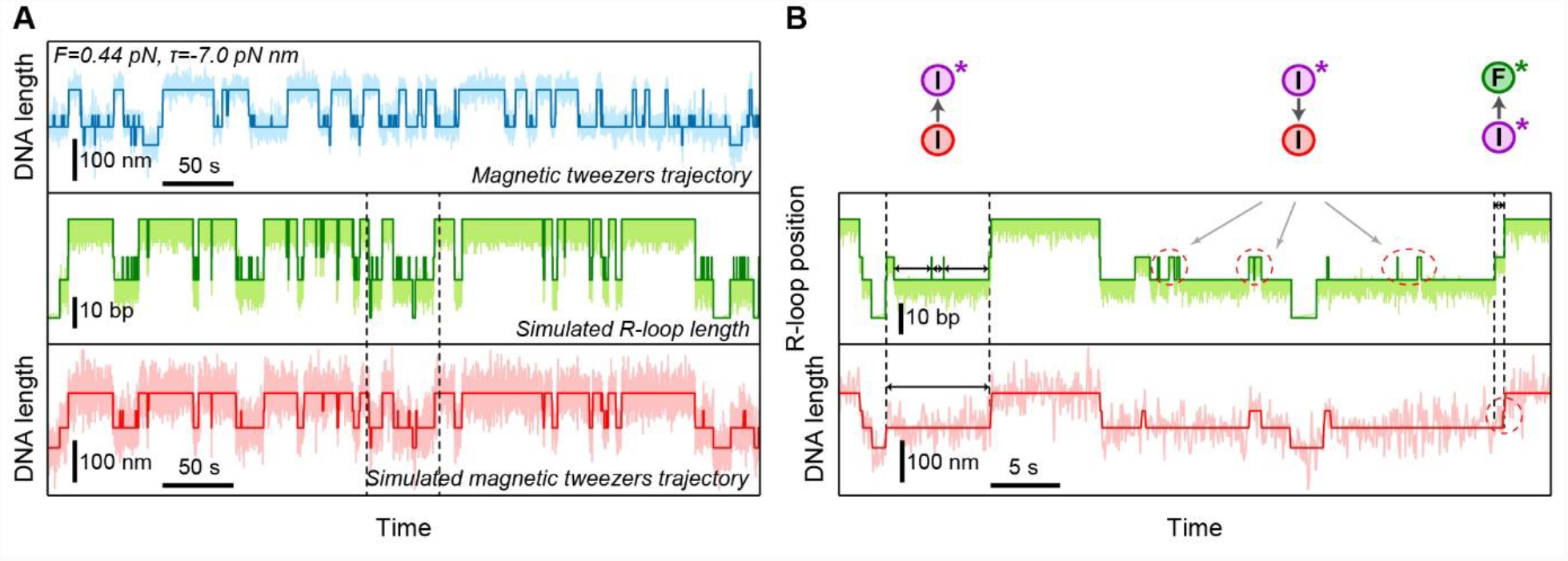
Simulations of R-loop length and magnetic tweezers experiments for double mismatch targets. Related to Figure 5. (A) Comparison between trajectories of a magnetic tweezers experiment (blue), a random walk simulation of the R-loop length (green) and a Brownian dynamics simulation of a magnetic tweezers experiment (red) for a target containing C:C mismatches at the positions 11 and 17 and 6 terminal mismatches. Light colors show the raw results of the experiments and simulations while dark colors show corresponding 3-state approximations. (B) Enlarged view into the simulated trajectories from areas separated by dashed lines in (A) depicting transitions extracted by the 4-state approximations of the simulated magnetic tweezers trajectories. Dashed areas depict transitions that are affected by measurement limitations.

**Figure S6.**
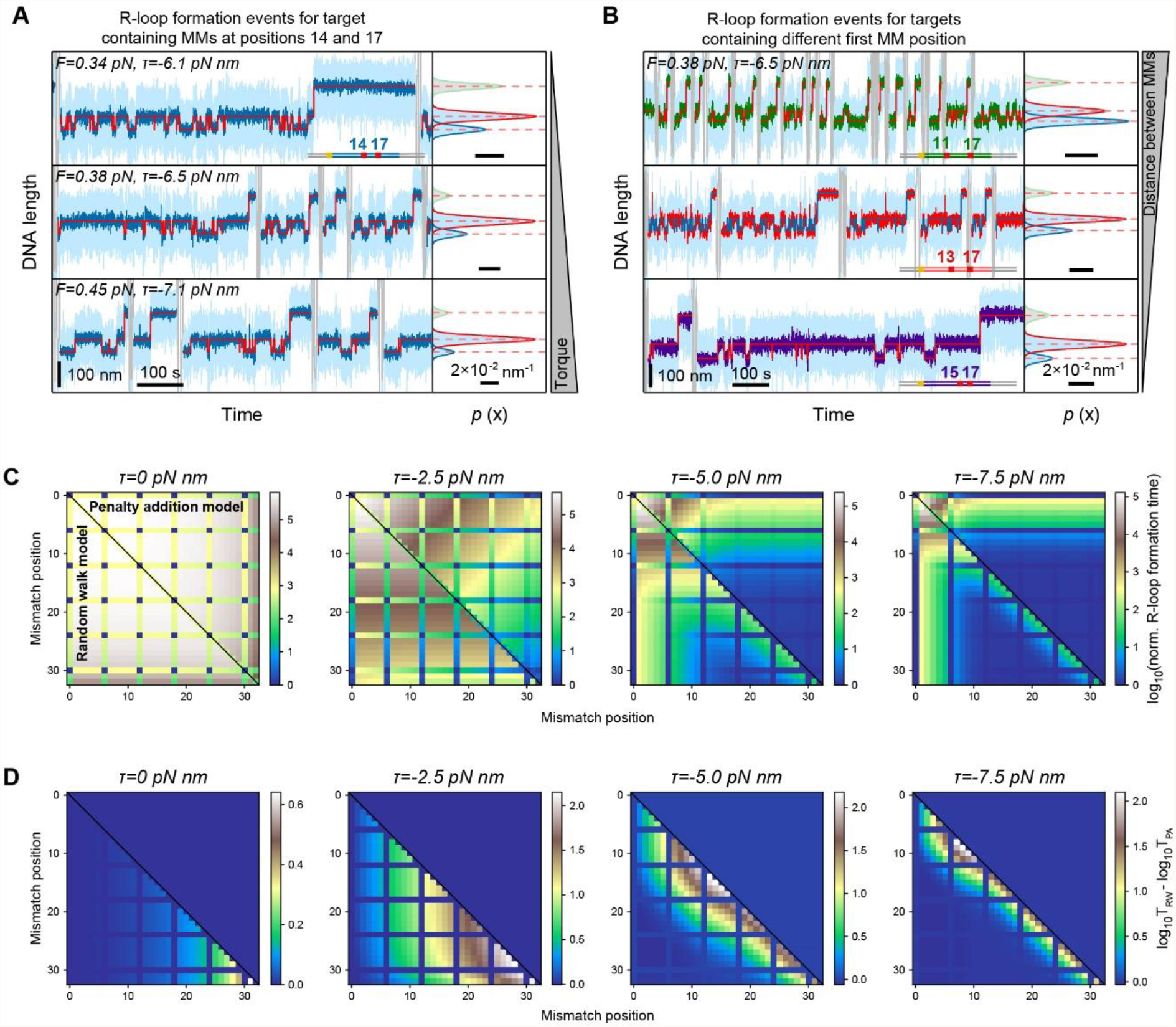
Impact of double mismatches on the R-loop formation times and transition rates between the states. Related to Figure 6. (A) Successive R-loop formation events measured at different torques (as indicated) for mismatches at position 14 and 17. Blue trajectory sections represent the actual R-loop formation events, while gray sections correspond to changes of the DNA supercoiling. The width of a formation event is roughly proportional to the R-loop formation time, such that a visual impression of the R-loop formation time for the different mismatches and torques can be obtained. The panels on the right side show histograms of the DNA length with a triple Gaussian fit to the U, I and F states. Due to the low occupancy the I* state was omitted. Light blue trajectories represent raw DNA length data collected at 120 Hz, while dark trajectories are after data filtering to 3 Hz. (B) Successive R-loop formation events for mismatches at varying first mismatch position, depicted as in (A). Blue trajectory represents the R-loop formation events for targets containing mismatches at positions 11 and 17, red – 13 and 17, violet – 15 and 17. (C) Normalized R-loop formation times for the different position combinations of double mismatches calculated for the random walk model (lower half) and for a simple addition of energy penalties (upper half) at different torques. (D) Difference between the R-loop formation times of the random walk model (T_RW_) and the simple penalty addition (T_PA_) calculated by normalizing the plots in C by the results from simple penalty addition.

**Table S1.** Best fit parameters for Figure 3 experiment. Related to Figure 3.

| Mismatch type | Mismatch position | $k_{step}$ ($s^{-1}$) | $\Delta G_{ini}$ ($k_B T$) | $\Delta G_{MM}$ ($k_B T$) | $\Delta G_{MM}$ ($k_B T$)*** |
| --- | --- | --- | --- | --- | --- |
| C:C* | 11 | 2000** | - | 7.6±0.4 | 9.2 |
|  | 13 | 3300±600 | 9.4±0.3 | 7.8±0.6 | 9.2 |
|  | 14 | 1800±300 | 9.6±0.2 | 7.4±0.4 | 9.2 |
|  | 15 | 4100±600 | 10.4±0.5 | 7.3±0.4 | 9.2 |
|  | 16 | 2500±1200 | 10.8±1.6 | 8.1±0.6 | 9.2 |
|  | 17 | 2100±100 | 11.2±0.5 | 8.0±0.2 | 9.2 |
| C:C | 17 | 640±70 | 8.8±0.5 | 6.7±0.3 | 10.5 |
| C:T | 17 | 630±50 | 8.7±0.6 | 4.3±0.2 | 8.3 |
| C:A | 17 | 760±70 | 9.3±0.5 | 2.7±0.1 | 7.5 |

\* Target sequences used here are designed to have the same neighboring bases and their sequences are presented in Table S3.
\*\* $k_{step}$ and $\Delta G_{MM}$ for C:C mismatch was obtained only from the fitting of $k_3$ .
\*\*\* Determined for DNA:DNA duplex from nucleic acid thermodynamics using NUPAC (Zadeh et al., 2011).

**Table S2.**
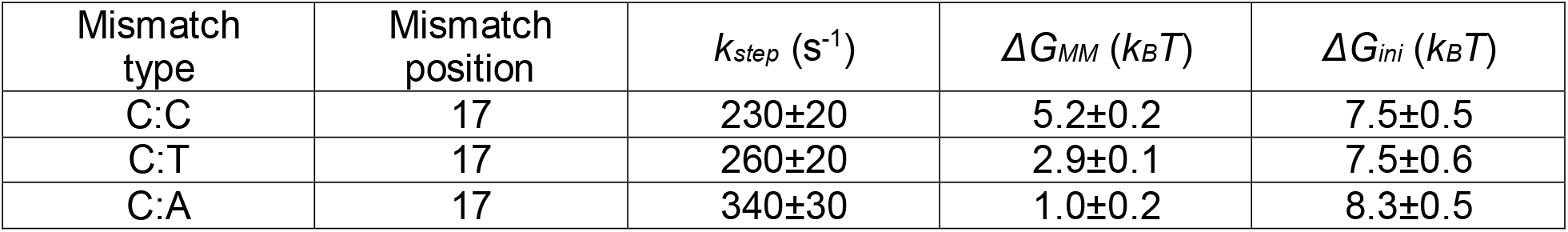
Best fit parameters for Figure 3 experiments when including the prebias from Figure 4 experiments. Related to Figure 3.

**Table S3.**
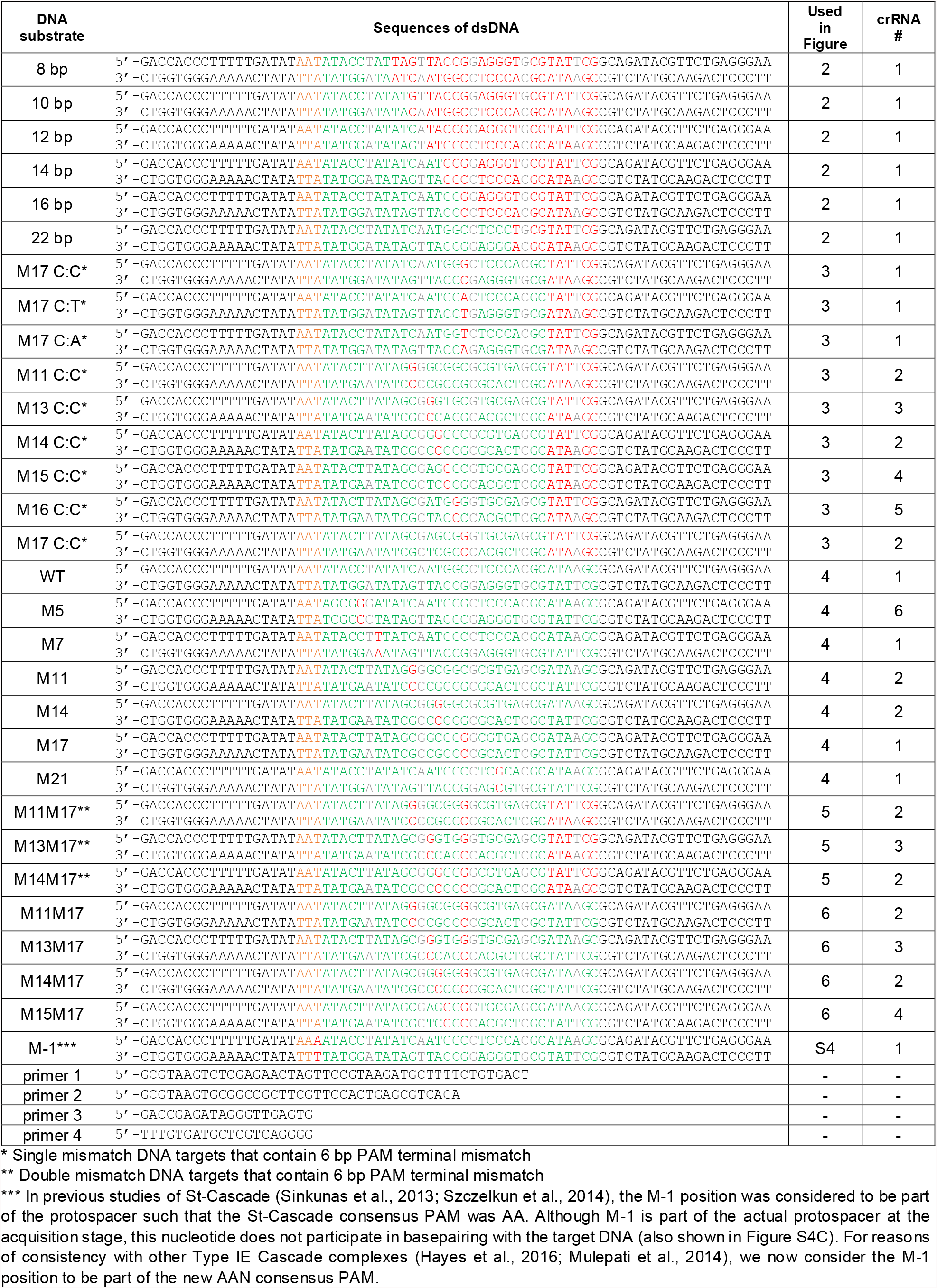
Sequences of the oligonucleotides used in this study to produce target DNAs. PAM*** for Cascade is coloured yellow, matching part of the target sequence – green, mismatched bases of the target sequence – red, flipped out bases – grey. Related to Figures 2-6.

**Table S4.**
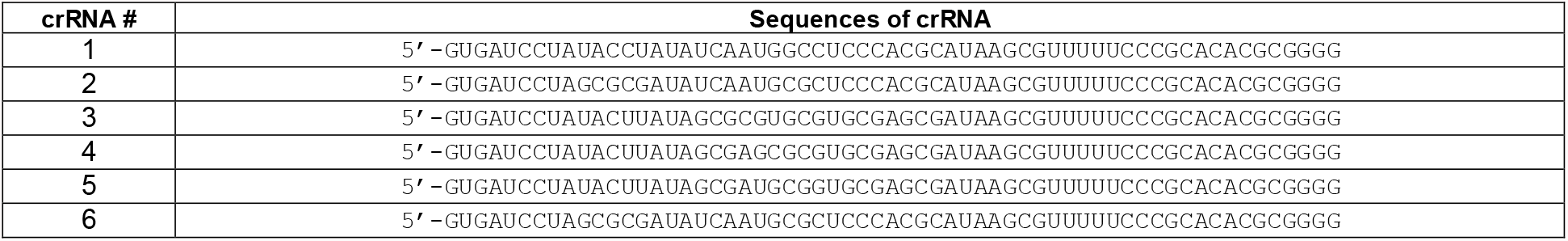
Sequences of crRNAs present in Cascade complexes used in this study. Related to Figures 2-6.

**Table S5.**
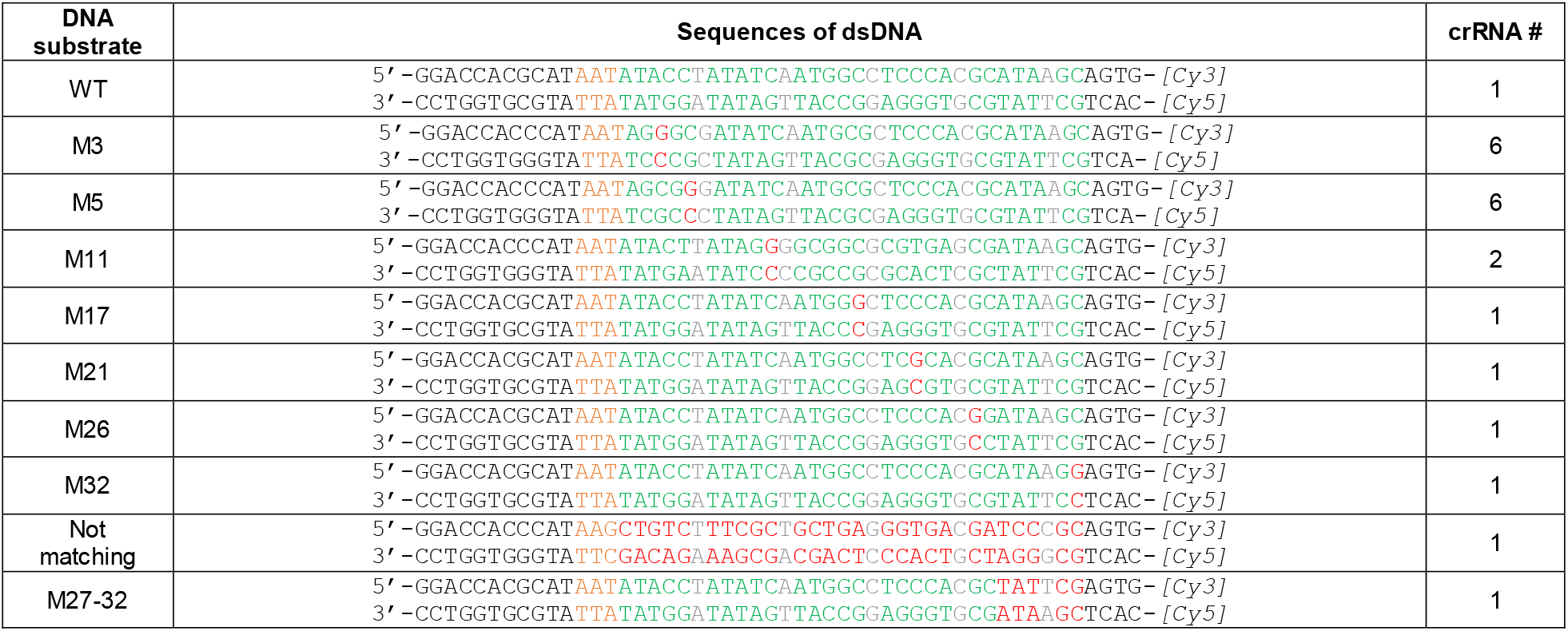
Oligonucleotide sequences used for the fluorescence measurements. PAM for Cascade is coloured yellow, matching part of the target sequence – green, mismatched bases of the target sequence – red, flipped out bases – grey. Related to Figure 4.

